# SLAP2 adaptor binding disrupts c-CBL autoinhibition to activate ubiquitin ligase function

**DOI:** 10.1101/2020.06.18.159806

**Authors:** Leanne E. Wybenga-Groot, Andrea J. Tench, Craig D. Simpson, Jonathan St. Germain, Brian Raught, Michael F. Moran, C. Jane McGlade

**Affiliations:** The Arthur and Sonia Labatt Brain Tumour Research Centre, The Hospital for Sick Children, 555 University Avenue, Toronto, ON, Canada, M5G 1X8; Program in Cell Biology, The Hospital for Sick Children, 555 University Avenue, Toronto, ON, Canada, M5G 1X8; SPARC BioCentre, The Hospital for Sick Children, 555 University Avenue, Toronto, ON, Canada, M5G 1X8; Department of Medical Biophysics, University of Toronto, 610 University Avenue, Toronto, ON, Canada, M5G 2M9; Princess Margaret Cancer Centre, University Health Network, Toronto, ON, M5G 1L7; Department of Molecular Genetics, 1 King’s College Circle, Toronto ON M5S 1A8

**Keywords:** Src-like adaptor protein, CBL, protein-protein interaction, ubiquitin ligase, X-ray crystal structure, activation mechanism

## Abstract

CBL is a RING type E3 ubiquitin ligase that functions as a negative regulator of tyrosine kinase signaling and loss of CBL E3 function is implicated in several forms of leukemia. The Src-like adaptor proteins (SLAP/SLAP2) bind to CBL and are required for CBL-dependent downregulation of antigen receptor, cytokine receptor, and receptor tyrosine kinase signaling. Despite the established role of SLAP/SLAP2 in regulating CBL activity, the nature of the interaction and the mechanisms involved are not known. To understand the molecular basis of the interaction between SLAP/SLAP2 and CBL, we solved the crystal structure of CBL tyrosine kinase binding domain (TKBD) in complex with SLAP2. The carboxy-terminal region of SLAP2 adopts an α-helical structure which binds in a cleft between the 4H, EF-hand, and SH2 domains of the TKBD. This SLAP2 binding site is remote from the canonical TKBD phospho-tyrosine peptide binding site but overlaps with a region important for stabilizing CBL in its autoinhibited conformation. In addition, binding of SLAP2 to CBL *in vitro* activates the ubiquitin ligase function of autoinhibited CBL. Disruption of the CBL/SLAP2 interface through mutagenesis demonstrated a role for this protein-protein interaction in regulation of CBL E3 ligase activity in cells. Our results reveal that SLAP2 binding to a regulatory cleft of the TKBD provides an alternative mechanism for activation of CBL ubiquitin ligase function.

## Introduction

Ubiquitin (Ub) modification of proteins regulates cellular processes including cell cycle, DNA repair, endocytosis, and signal transduction. The ubiquitination machinery consists of a cascade of E1, E2, and E3 components, which activate and transfer ubiquitin to substrates in a sequential manner (Hershko & Ciechanover, 1998, Kerscher, Felberbaum et al., 2006). Substrate specificity in the ubiquitin system is determined through a large and diverse family of E3 ubiquitin ligases that bind target proteins (Deshaies & Joazeiro, 2009, Rotin & Kumar, 2009). The Casitas B-cell lymphoma (*CBL*) gene encodes the E3 ubiquitin ligase CBL (also referred to as c-CBL), a central regulator of tyrosine kinase (TK) signaling (Clague, Liu et al., 2012, Joazeiro, Wing et al., 1999, Marmor & Yarden, 2004, Mohapatra, Ahmad et al., 2013, Thien & Langdon, 2005). Ubiquitination of activated receptor TKs by CBL nucleates the assembly of endocytic proteins both at the membrane and at sorting endosomes to promote lysosome targeting, degradation and signal termination (Acconcia, Sigismund et al., 2009, Urbe, 2005). CBL is also important for down regulation of signaling from antigen and cytokine receptors through ubiquitination of receptor chains and associated cytosolic TKs, leading to inactivation and/or proteosomal degradation (Andoniou, Lill et al., 2000, Thien, Bowtell et al., 1999, Yokouchi, Kondo et al., 2001).

CBL consists of an amino-terminal tyrosine kinase binding domain (TKBD), a linker helix region (LHR) and a really interesting new gene (RING) domain, followed by a carboxy-terminal region containing binding sites for Src homology 2 (SH2) and Src homology 3 (SH3) domain containing signaling and adaptor proteins(Mohapatra et al., 2013). CBL TKBD is composed of a four-helix bundle (4H), an EF-hand, and a variant SH2 domain, which binds substrates, such as activated TKs, in a phospho-tyrosine dependent manner(Meng, Sawasdikosol et al., 1999). The CBL RING domain binds E2 conjugating enzymes required for ubiquitin transfer to TKBD bound substrates(Joazeiro et al., 1999, Levkowitz, Waterman et al., 1999, Ryan, Sivadasan-Nair et al., 2010). Recruitment of CBL to activated TK complexes leads to phosphorylation at a conserved tyrosine residue (Tyr371) within the CBL LHR, and activation of its E3 ligase function (Kassenbrock & Anderson, 2004, Levkowitz et al., 1999, Liu, Kimura et al., 2002). Structural studies have shown that unphosphorylated CBL adopts a closed autoinhibited conformation with Tyr371 forming part of a LHR-TKBD interface (Dou, Buetow et al., 2012, Zheng, Wang et al., 2000). Phosphorylation of Tyr371 releases the LHR-TKBD interaction causing a conformational change that places the E2 active site in close proximity to TKBD bound substrate, and stabilizing E2 bound ubiquitin for transfer (Dou et al., 2012, Dou, Buetow et al., 2013). Missense mutations that impair this E3 ligase activation mechanism are implicated in several forms of myeloid leukemia and highlight the importance of CBL mediated ubiquitination in the regulation of TK signaling (Dou et al., 2012, Kales, Ryan et al., 2010, Loh, Sakai et al., 2009, Ogawa, Shih et al., 2010).

Src-like adaptor proteins (SLAP and SLAP2) play important roles in CBL mediated down regulation of antigen receptor, cytokine receptor, and RTK signaling (Dragone, Myers et al., 2006a, Dragone, Shaw et al., 2009, Lebigot, Gardellin et al., 2003, Loreto, Berry et al., 2002, Loreto & McGlade, 2003, Myers, Sosinowski et al., 2006, Naramura, Kole et al., 1998, Sosinowski, Killeen et al., 2001, Sosinowski, Pandey et al., 2000). SLAP and SLAP2, which are membrane associated by an amino-terminal myristoylation site, contain adjacent SH3 and SH2 domains that associate with one another through β-sheet formation, followed by a carboxy (C)-terminal tail region lacking obvious domains or protein interaction motifs (Loreto et al., 2002, Pandey, Duan et al., 1995, Pawson, 2007, Tang, Sawasdikosol et al., 1999, Wybenga-Groot & McGlade, 2013, Wybenga-Groot & McGlade, 2015). Recruitment of SLAP/SLAP2 to activated receptor TKs is mediated through the SH3/SH2 module, while the C-terminal tail region of both SLAP and SLAP2 constitutively associates with CBL (Holland, Liao et al., 2001, Loreto & McGlade, 2003, Myers et al., 2006). SLAP association with CBL is important for ubiquitination of substrates ZAP-70 and TCRξ, as well as RTKs including FLT3 and c-KIT though the mechanisms involved are unknown (Abe, Kiyoi et al., 2006, Kazi, Agarwal et al., 2014, Kazi & Ronnstrand, 2012, Liontos, Dissanayake et al., 2011, Loreto et al., 2002, Myers et al., 2006, Sosinowski et al., 2000, Wybenga-Groot & McGlade, 2015).

CBL binds to multiple SH2 and SH3 domain containing adaptor proteins through canonical binding motifs found in the CBL carboxy-terminal region while SLAP and SLAP2 binding requires a region of the SLAP/SLAP2 C-terminal tail and the CBL TKBD (Loreto et al., 2002, Pandey et al., 1995, Tang et al., 1999). SLAP/SLAP2 binding to CBL is also distinct from other TKBD binding proteins as it is independent of tyrosine phosphorylation and is not disrupted by a CBL Gly306Glu mutation, which abolishes TKBD binding to tyrosine phosphorylated substrates (Loreto et al., 2002). To understand the molecular basis of this interaction, we solved the X-ray crystal structure of CBL TKBD in complex with SLAP2. We describe how a C-terminal region of SLAP2 adopts an α-helical structure that binds CBL TKBD in the same cleft that is occupied by the LHR in the autoinhibited CBL conformation. In this way, SLAP2 binding precludes CBL autoinhibition, and stimulates CBL autoubiquitination activity. This indicates that in addition to its adaptor functions, SLAP/SLAP2 binding contributes to CBL activation.

## Results

### Crystal structure of CBL TKBD in complex with SLAP2 defines a novel binding interface

SLAP2 interacts with the CBL TKBD via a phosphorylation independent mechanism involving its C-terminal region (Fig. 1A) (Holland et al., 2001, Loreto et al., 2002, Manes, Masendycz et al., 2006). To understand the molecular basis of this interaction, we determined the crystal structure of CBL TKBD in complex with a portion of the SLAP2 C-terminal tail (Figure 1B, Table 1). The crystal asymmetric unit contains two CBL TKBD molecules and two SLAP2 molecules (residues 237-255) related by a two-fold symmetry axis to form a CBL/SLAP2 dimer (Fig. 1B). The CBL TKBD structure is well ordered except for the N-terminal (25-47 molecule 1, 25-51 molecule 2) and C-terminal (353-357 molecule 1, 352-357 molecule 2) residues. Superposition of CBL/SLAP2 with native CBL TKBD (PDB id: 1B47) indicates that CBL adopts the typical integrated module, composed of a calcium-bound EF-hand wedged between a four-helix bundle (4H) and a divergent SH2 domain, with root mean square deviation (rmsd) of 0.9 Å(main chain atoms) for CBL TKBD (Fig. 1C)(Meng et al., 1999). Superposition of the 4H bundle and EF-hand of CBL/SLAP2 with those of native and liganded CBL TKBD (PDB id: 2CBL) shows that binding of the C-terminal portion of SLAP2 to CBL does not induce closure of the domains to the extent that phosphopeptide binding to the SH2 domain does, such that CBL/SLAP2 most resembles unliganded CBL TKBD (Fig. 1C) (Meng et al., 1999).

**Table 1.**
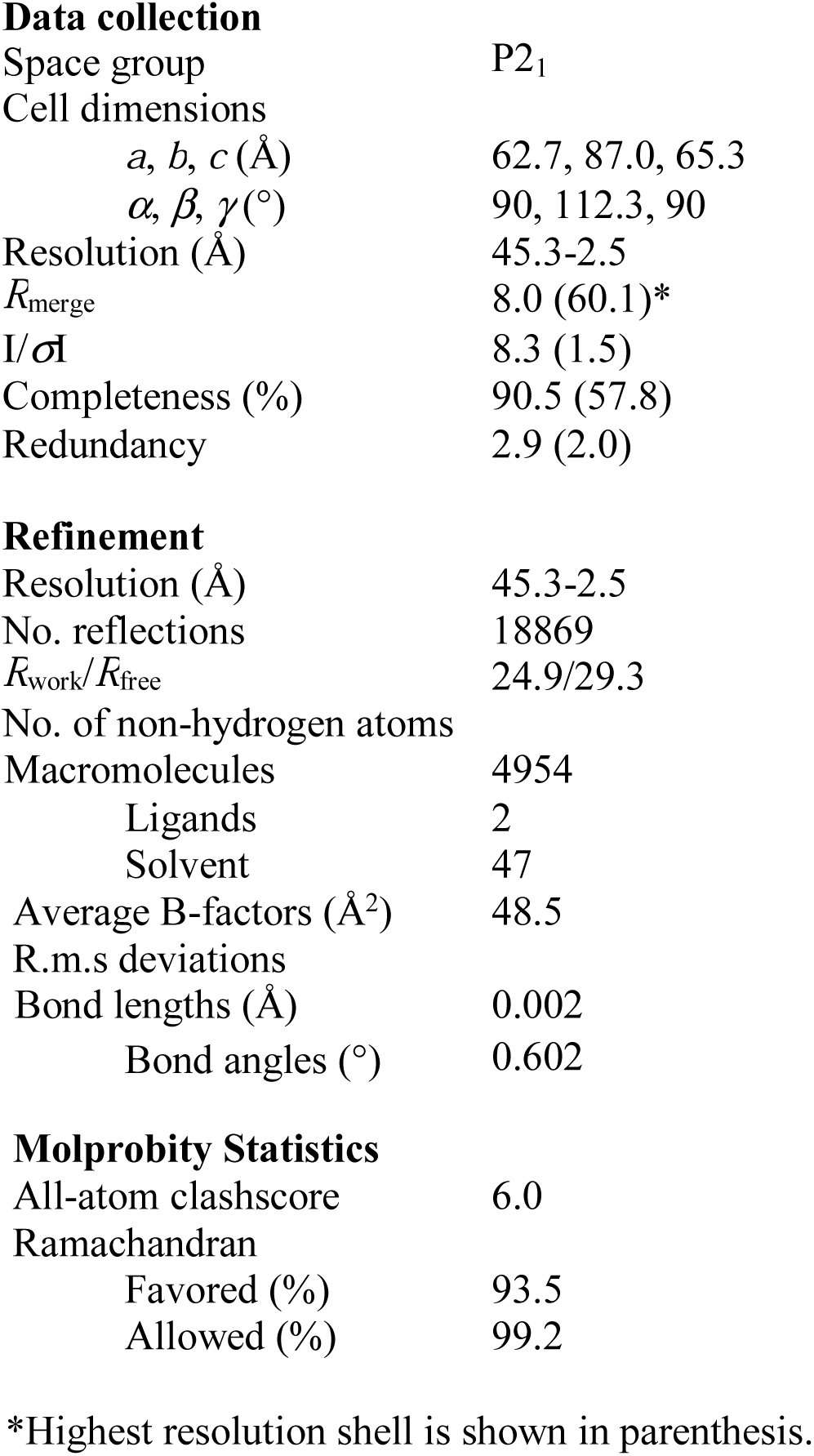
Data collection and refinement statistics

**Figure 1:**
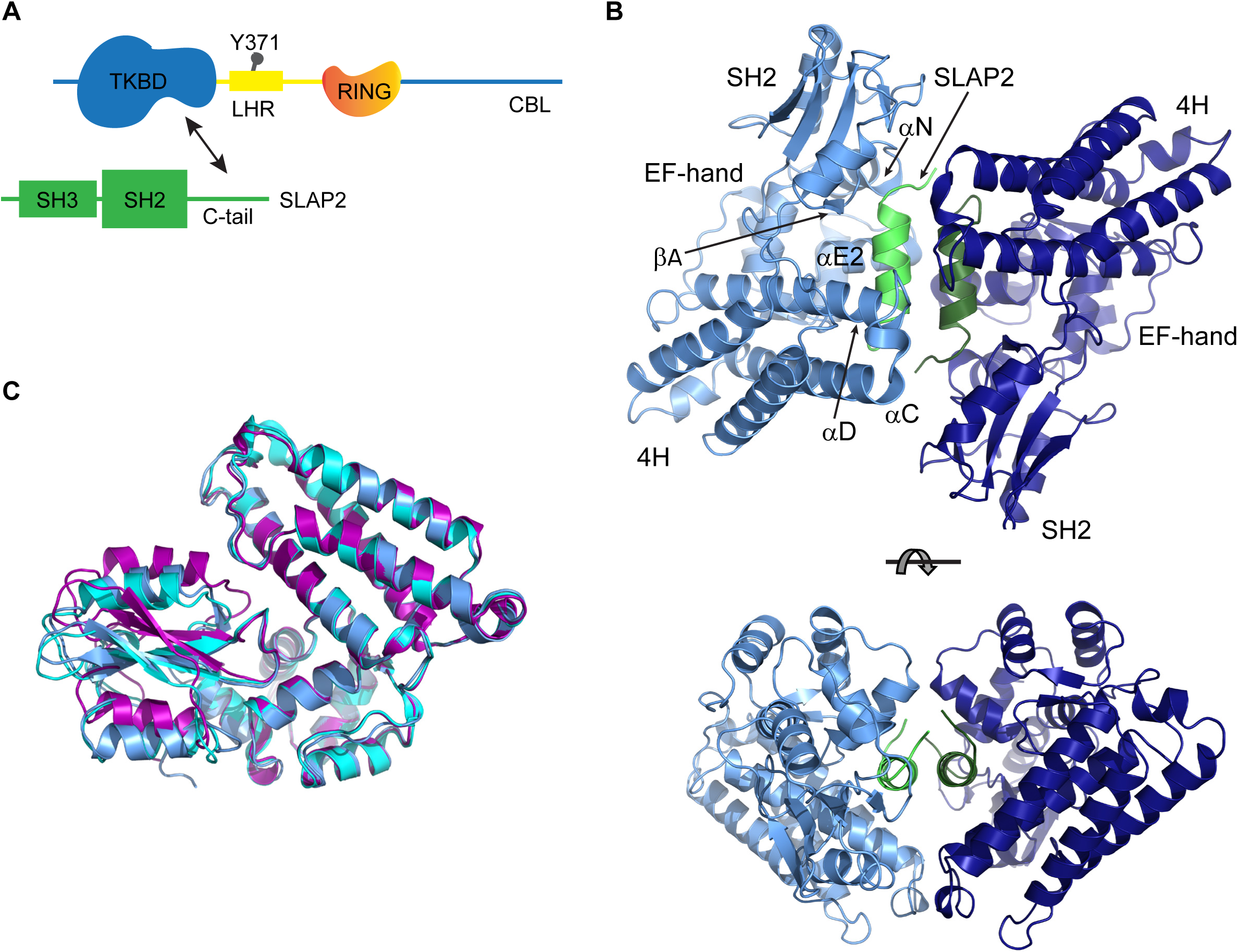
Crystal Structure of CBL TKBD with C-terminal tail of SLAP2. A) Cartoon of CBL and SLAP2 domains, with arrow indicating interaction between TKBD and SLAP2 C-tail region. B) Ribbon representations of the Cα atoms of the CBL/SLAP2 crystal structure, with CBL monomers in blue and SLAP2 monomers in green and elements of CBL TKBD labelled. Upper and lower panels are related by a rotation of approximately 90° about the horizontal axis. C) Superposition of the Cα atoms of the 4H bundle and EF-hand of CBL/SLAP2 (PDB id: 6XAR) with those of unliganded (PDB id: 1B47) and liganded CBL TKBD (PDB id: 2Cbl), shown in blue, cyan, and magenta, respectively. All ribbon diagrams were prepared with Pymol (The PyMOL Molecular Graphics System, Version 1.5.0.4 Schrödinger, LLC.).

The SLAP2 C-terminal tail binds as an α-helix in a cleft formed by the three subdomains of TKBD, opposite the phospho-tyrosine peptide binding pocket (Fig. 2A). The SLAP2 binding cleft is framed by helices αC and αD of the 4H bundle, helix αE2 and loop αE2-αF2 of the EF-hand, and helix αN, loop αN-βA, and strand βA of the SH2 domain (Fig. 2A,B) (Meng et al., 1999). The CBL/SLAP2 interface is stabilized by hydrophobic interactions involving side chains from Leu241, Leu245, and Leu251 of SLAP2 and Lys153, Leu154, Met222, Ala223, Trp258, and Val263 of CBL, and by hydrogen bonds involving both backbone carbonyl groups and side chain atoms (see Supplementary Table 1 and Supplementary Fig. 1). Altogether, the interaction between CBL and SLAP2 results in 4592 Å^2^ of overall contact area. The SLAP2 α-helices interact expansively, stabilized by extensive hydrogen bonding and hydrophobic interactions, while few interactions are formed between the CBL monomers (Supplementary Fig. 2 and Supplementary Tables 2 and 3).

**Figure 2:**
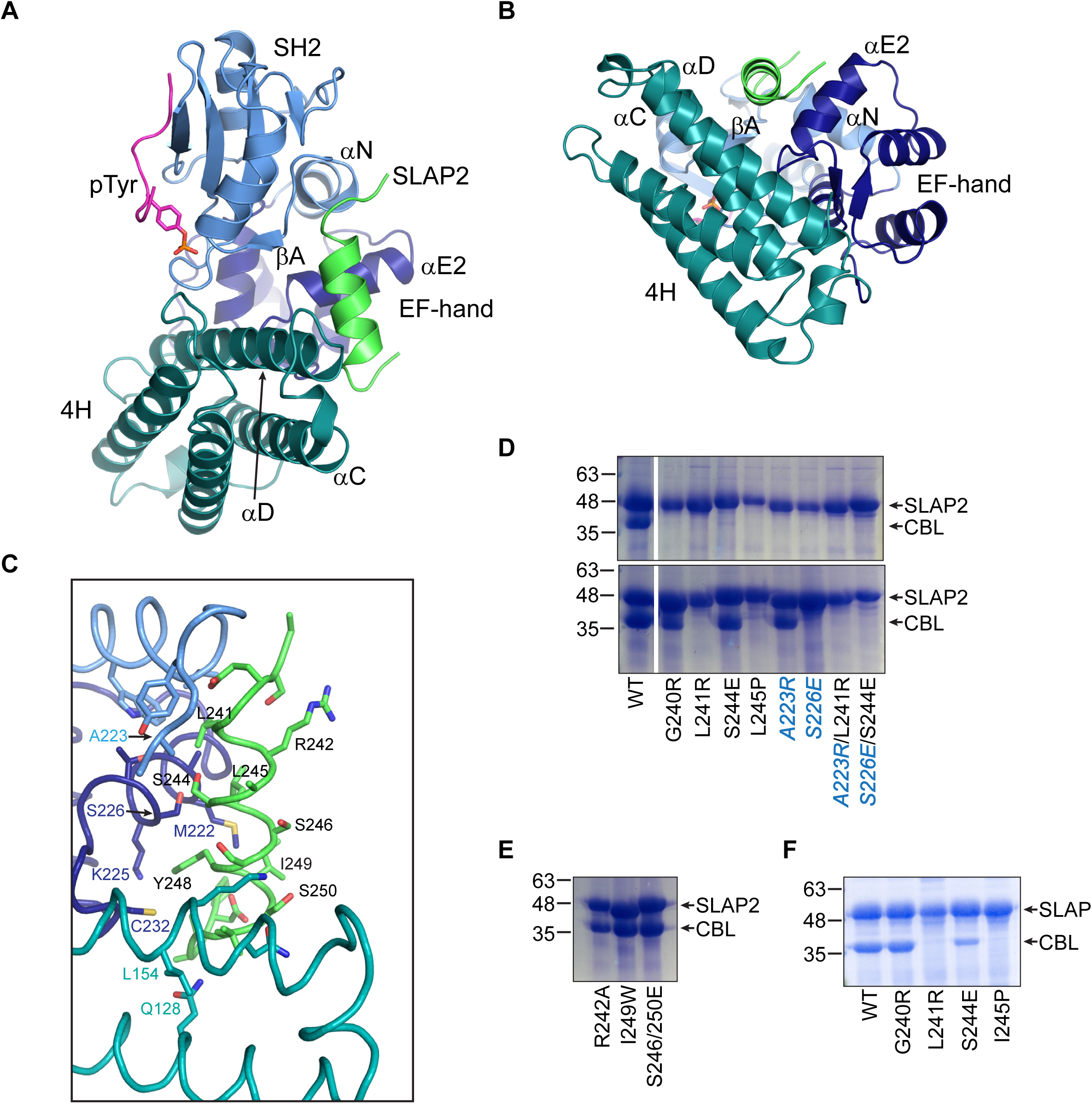
Validation of CBL/SLAP2 interaction by mutagenesis. A) Ribbon representation of the Cα atoms of the CBL/SLAP2 crystal structure, with a pTyr peptide modeled in magenta. Elements of CBL TKBD are labelled, with the 4H bundle shown in teal, EF-hand in dark blue, and SH2 domain in light blue. SLAP2 is coloured green. B) Ribbon representation of the Cα atoms of CBL/SLAP2 structure coloured as in A, highlighting the CBL regulatory cleft. C) Magnification of the CBL/SLAP2 interface, coloured as in A, with side chains of CBL and SLAP2 molecule shown in stick and labelled in teal/dark blue/light blue according to their respective subdomains for CBL and in black for SLAP2. Oxygen, nitrogen, and sulfur atoms are coloured red, blue, and yellow, respectively. D) SDS-PAGE gel stained with Coomassie blue for CBL and SLAP2 proteins co-purified under high salt (upper panel) and low salt (lower panel) conditions from Duet-His-Δlinker-hSLAP2-CBL and its mutated variants. SLAP2 and CBL mutants are labelled in black and blue, respectively. E) As in D, for SLAP2/SLAP2 interface mutants, co-purified in low salt conditions. F) As in D, for CBL and SLAP proteins co-purified in low salt conditions from Duet-His-mSLAP-CBL and its mutated variants.

To validate the CBL/SLAP2 binding interface, a series of mutants in a Duet co-expression vector system was generated for co-purification trials. The SLAP2 sequence was preceded by thioredoxin (Trx) and His-tag fusion proteins to allow co-purification of CBL TKBD that was bound to Trx-His-SLAP2 during the initial affinity chromatography purification step. Surface residues deemed to be important to the CBL/SLAP2 interface, but not CBL function or folding, were mutated to disrupt CBL/SLAP2 binding (Fig. 2C and Supplementary Fig. 1). In our structure, SLAP2 Gly240 locates close to CBL Met269 and Ala270, such that mutation to a bulky Arg residue (G240R) could cause sterical constraints. Similarly, SLAP2 Leu241 is situated in a hydrophobic cleft composed of CBL Ala223, Trp258, and Val263; mutation to Arg (L241R) was predicted to disturb this binding pocket. Mutation of CBL Ala223 to Arg (*A223R*) would likewise disrupt this pocket, with CBL/SLAP2 double mutant *A223R*/L241R predicted to drastically disturb binding through sterical constraints and charge repulsion. (For clarity, CBL mutations herein are indicated in italics, SLAP2 in regular font.) Similarly, single or double mutation of CBL Ser226 or SLAP2 Ser244, whose side chain hydroxyl groups are in close proximity to one another, to charged Glu residues (*S226E* and S244E) was anticipated to interfere with CBL/SLAP2 binding. Finally, given that SLAP2 binds CBL as an α-helix, mutation of Leu245, located in the center of this α-helix, to proline (L245P) was expected to interrupt the helical structure and thus prevent SLAP2 binding.

CBL TKBD co-purified with SLAP2 in high salt conditions (0.5 M NaCl), while mutation of either SLAP2 or CBL at interface residues resulted in loss of CBL co-purification (Fig. 2D, upper panel). In less stringent low salt concentration (0.15 M NaCl) mutations at L241R, L245P, and *S226E* still abrogated CBL co-purification, while G240R, S244E, and *A223R* mutations were tolerated, resulting in CBL co-purification (Fig. 2D, lower panel). However, double mutation of the CBL/SLAP2 interface (*A223R*/L241R and *S226E*/S244E; Fig. 2D, lower panel) was not tolerated, even at low salt concentration. In contrast, mutation of SLAP2 residues involved in the SLAP2/SLAP2 interface but not CBL binding (R242A, I249W, S246/250E) did not significantly impact co-purification of CBL, suggesting that TKBD binding does not require a SLAP2 dimer (Fig. 2E).

The related adaptor protein SLAP also interacts with CBL TKBD (Tang et al., 1999), and its amino acid sequence corresponding to the SLAP2 C-terminal α-helical region is highly conserved (Fig. 3A). To determine if SLAP binds CBL TKBD by a similar mechanism, we tested SLAP and a set of analogous interface mutants in co-purification experiments. Like SLAP2, SLAP mutations at L241R and I245P abrogated CBL co-purification, while mutations G240R and S224E were tolerated (Fig. 2F). Together, these studies indicate that SLAP and SLAP2 interact with CBL via an α-helix near their C-terminus, which binds to a cleft of the CBL TKBD that is distinct from the canonical phospho-tyrosine binding site in the CBL SH2 domain.

**Figure 3:**
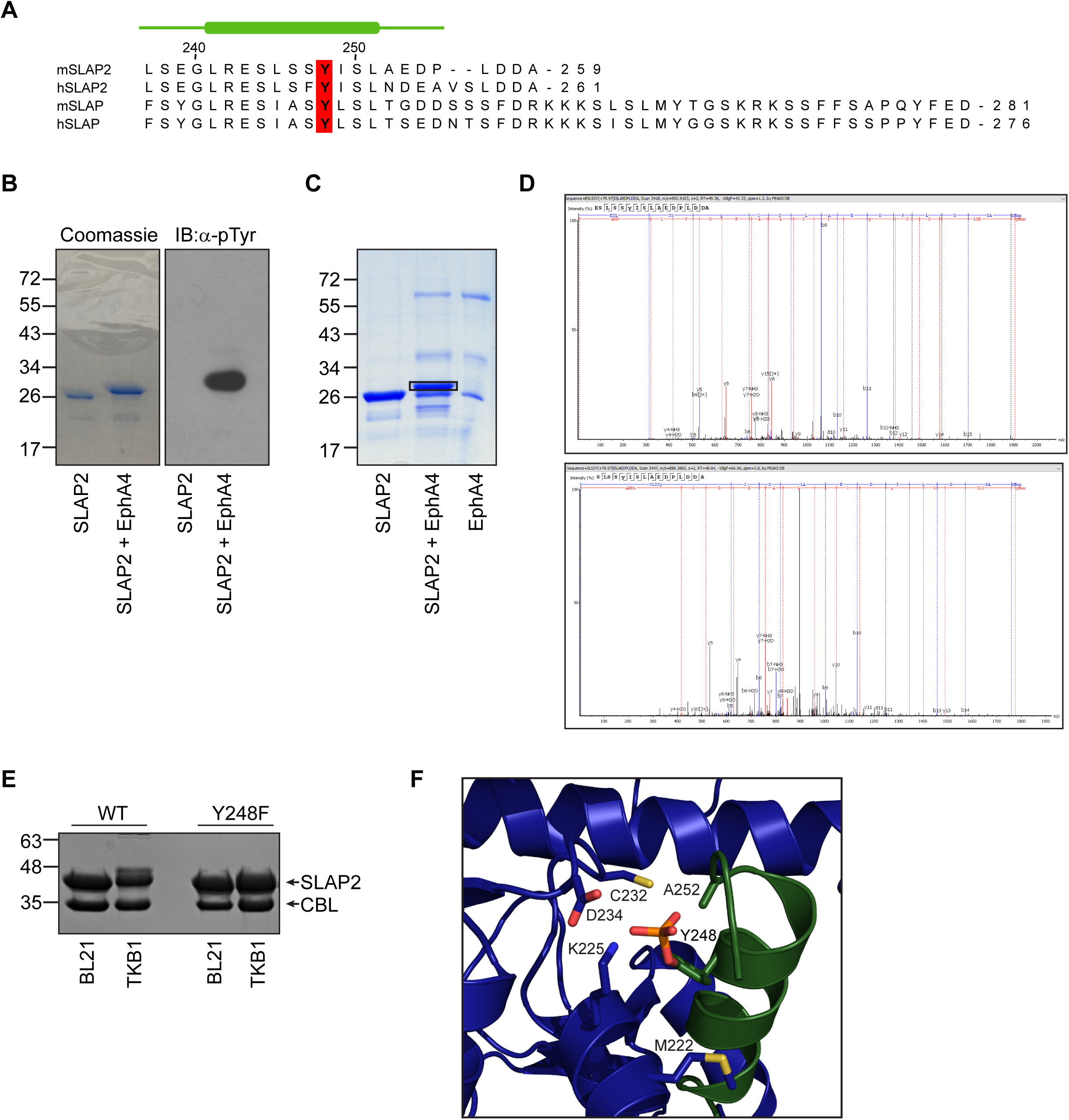
Phosphorylation of SLAP2 at tyrosine 248. A) Sequence alignment of the C-tail region of SLAP proteins, with residue numbering based on human SLAP2. Conserved Tyr248 is marked with a red box. The secondary structure elements of mSLAP2 as determined by X-ray crystallography are shown in green, with the line and box representing coil and α-helix, respectively. B) SDS-PAGE gel stained with Coomassie blue for SLAP2 and SLAP2 incubated with EphA4 kinase under phosphorylating conditions (left panel). Immunoblot of SLAP2 +/- EphA4, immunoblotted with anti-pTyr (right panel). C) SDS-PAGE gel as in A, with EphA4 alone also shown. Box indicates upper gel band excised for mass spectrometry analysis. D) Intensity versus charge to mass ratio (m/z) plot for SLAP2 C-tail peptides ESLSSY_248_ISLAEDPLDDA (upper panel) and SLSSY_248_ISLAEDPLDDA (lower panel) generated by protease digestion of gel band from B with trypsin and GluC, respectively, followed by LC-MS/MS analysis. E) SDS-PAGE gel stained with Coomassie blue for CBL and SLAP2 WT or Y248F co-purified under low salt conditions from Duet-His-Δlinker-hSLAP2-CBL expressed in normal (BL21) or phosphorylating (TKB1) conditions. F) Ribbon representation of the Cα atoms of the CBL/SLAP2 interface, with CBL shown in blue and SLAP2 in green. Residues of CBL and SLAP2 are shown in stick with carbon atoms coloured according to their respective backbones, and oxygen, nitrogen, sulfur, and phosphorus atoms coloured red, blue, yellow, and orange, respectively. A phosphate group has been modeled on the side chain hydroxyl group of tyrosine 248.

Further examination of the SLAP/SLAP2 α-helix protein sequence revealed a conserved tyrosine residue (Tyr248) (Fig. 3A) that is predicted to be phosphorylated by the Kinexus phosphorylation site prediction algorithm, but to our knowledge has not been experimentally confirmed (www.phosphonet.ca). SLAP2 has been shown to be tyrosine phosphorylated in CSF-1 stimulated FD-Fms cells, although the site of phosphorylation was not determined(Pakuts, Debonneville et al., 2007). To identify SLAP2 tyrosine phosphorylation sites, purified SLAP2 protein was incubated *in vitro* with EphA4 kinase in the presence of ATP. Phosphorylated SLAP2 was detected by an upward shift in molecular weight by SDS-PAGE and confirmed by immunoblot analysis (Fig. 3B). Sites of phosphorylation were identified by in-gel protease digestion of the isolated, shifted band followed by liquid chromatography tandem mass spectrometry (LC-MS/MS), which identified Tyr248 as a site of phosphorylation (Fig. 3C,D). To investigate whether phosphorylation of Y248 impacts SLAP2 binding to CBL, co-purification experiments were performed under conditions where SLAP2 was tyrosine phosphorylated. CBL TKBD co-purified with phosphorylated SLAP2 WT as well as a Y248F mutant indicating that SLAP2 interaction with CBL is not disrupted by Y248 phosphorylation, nor is it required for binding (Fig. 3E). In agreement, a phosphate ion was modeled on the side chain hydroxyl of Tyr248 in the CBL/SLAP2 structure (Fig. 3F). The structure model accommodates phosphorylation at Tyr248 without causing sterical constraints at the CBL/SLAP2 interface.

### SLAP2 binding to CBL precludes inhibitory LHR-RING interactions with the TKBD

Unphosphorylated CBL adopts a closed, autoinhibited conformation in which the LHR and RING domain pack against the TKBD in a manner that restricts ubiquitin ligase activity (PDB id: 1FBV) (Fig. 4A) (Dou et al., 2012, Zheng et al., 2000). Phosphorylation of Tyr371 in the LHR disrupts these interactions, reorients the LHR-RING in proximity of substrates, and stabilizes E2 bound ubiquitin for transfer (Dou et al., 2012, Dou et al., 2013). Superposition of CBL/SLAP2 with autoinhibited CBL TKBD-LHR-RING plus E2 (PDB id:1FBV) revealed that SLAP2 binds to a similar region of CBL TKBD as the regulatory LHR and RING domain (Fig. 4B). The LHR forms an ordered loop and an α-helix that pack against the TKBD, with linker-TKDB interactions centered on linker residues Tyr368 and Tyr371(Zheng et al., 2000). In the CBL/SLAP2 structure, SLAP2 residue Leu241 occupies the same buried environment as Tyr371 in autoinhibited CBL (Fig. 4C). Moreover, side chain residues from helix αE2 of the EF-hand, such as Ala223 and Ser226, as well as Val263 from loop αN-βA of the SH2 domain, make multiple van der Waals contacts with the linker helix in autoinhibited CBL; these same residues are involved in the SLAP2 binding site (Fig. 2C, Supplementary Fig. 1, Supplementary Table 1). Similarly, the SLAP2 α-helix and its C-terminal extension occupy the same space as linker-loop 2 of the LHR and the N-terminus of the RING domain in closed CBL, respectively (Fig. 4C). Indeed, several residues which stabilize the TKBD-RING interface in closed CBL, including Gln128, Asn150, Lys153, and Leu154, are involved in SLAP2 binding in the CBL/SLAP2 structure (Supplementary Fig. 1, Supplementary Table 1). CBL residue Met222 stabilizes the TKBD-RING interface in autoinhibited CBL and forms part of an E2 binding pocket upon the addition of UbcH7 (Dou et al., 2012, Zheng et al., 2000). Notably, Met222 is also involved in hydrophobic interactions with Leu245 and Ile249 of the SLAP2 α-helix. These observations suggest a model in which SLAP2 binding to CBL would preclude LHR-RING interactions with the TKBD and favour the open, catalytically competent conformation (Fig. 4D).

**Figure 4:**
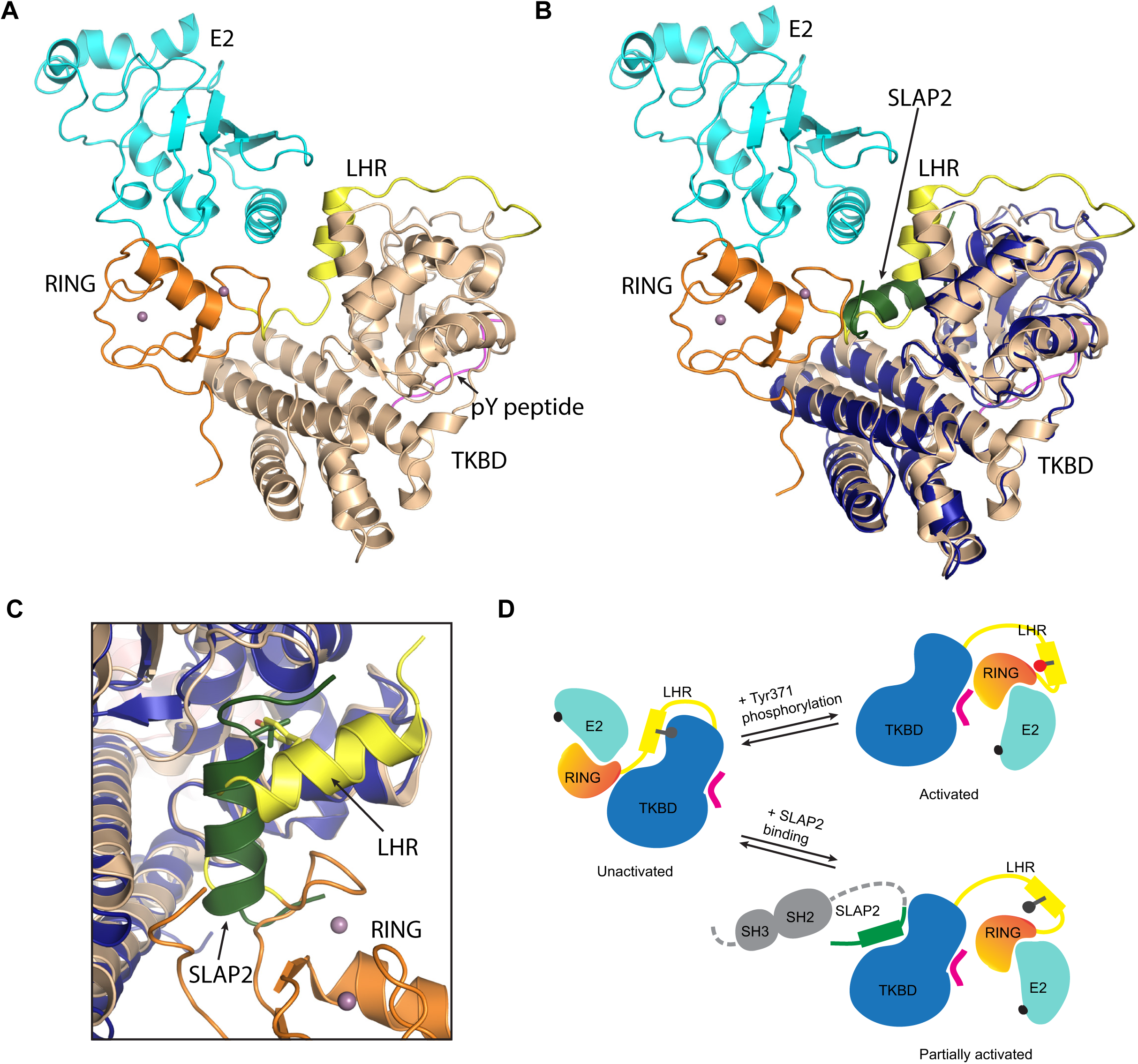
SLAP2 binding precludes LHR-RING interactions with the TKBD. A) Ribbon representation of the Cα atoms of the CBL TKBD-LHR-RING plus E2 (PDB id: 1FBV), with TKBD, LHR, RING, and E2 coloured beige, yellow, orange, and cyan, respectively. pTyr peptide and calcium ions are shown in magenta and mauve, respectively. B) Superposition of the Cα atoms of CBL/SLAP2 (PDB id: 6XAR) with CBL TKBD-LHR-RING plus E2 (PDB id: 1FBV), coloured as in A. CBL/SLAP2 is shown in dark blue and green. C) Magnification of the CBL regulatory cleft, with Leu241 of SLAP2 (green) and Tyr371 of CBL LHR (yellow) shown in stick (CBL LHR is from structure of autoinhibited CBL TKBD-LHR-RING, PDB id: 2Y1N). D) Model depicting CBL autoinhibition and conformational changes induced by phosphorylation of Y371 based on Dou *et al*. (Dou et al., 2012), and as a consequence of SLAP/SLAP2 binding. N-terminal regions of SLAP2 are shown in grey or hashed line, as their orientation with respect to CBL and the α-helical C-tail of SLAP2 are unknown.

### SLAP2 binding to CBL promotes ubiquitin ligase activity

Given that binding of SLAP2 and the LHR-RING would be mutually exclusive (Fig. 4C), we reasoned that binding of SLAP2 to CBL could displace the LHR-RING, thereby promoting the catalytically competent conformation (Fig. 4D). To test this hypothesis, an *in vitro* ubiquitination assay in which E3 ligase activity is detected by a smear at high molecular weight on an anti-ubiquitin (Ub) immunoblot was employed (Joazeiro et al., 1999, Kassenbrock & Anderson, 2004). As previously reported, purified CBL TKBD-LHR-RING (2-436) had no detectable ubiquitination activity, while its phosphorylated version (pCBL) displayed substantial activity, indicating the inactive and active forms of CBL, respectively (Fig. 5A) (Kassenbrock & Anderson, 2004). Addition of purified SLAP2 or phosphorylated SLAP2 (pSLAP2) to unphosphorylated CBL stimulated ubiquitination activity *in vitro* (Fig. 5A). Notably, addition of pSLAP2 to CBL appeared to promote CBL ubiquitination activity to a greater extent than addition of SLAP2 (Fig. 5A,B). Neither addition of SLAP2 nor pSLAP2 to active pCBL appeared to further enhance or diminish ubiquitination activity (Fig. 5A). To further compare and quantify CBL ubiquitin ligase activity, we employed a commercially available chemiluminescence based E3 ligase activity assay called E3LITE (Life Sensors), which measures the total amount of polyubiquitin chains formed in an *in vitro* reaction. In this assay, detection of polyubiquitylated protein increased upon addition of SLAP2 or pSLAP2 to CBL, compared to CBL alone (Fig. 5C and Supplementary Fig. 3A). Consistent with the anti-Ub immunoblot analysis, pSLAP2 stimulated CBL ligase activity to a greater extent than SLAP2 in the E3 ligase assay. Notably, the ubiquitin ligase activity of CBL/pSLAP2 remained less than that of pCBL in both activity assays (Fig. 5A,C). This is consistent with our model that pSLAP2 interaction with CBL is partially activating, such that a catalytically competent conformation is favoured upon SLAP2 binding (Fig. 4D).

**Figure 5:**
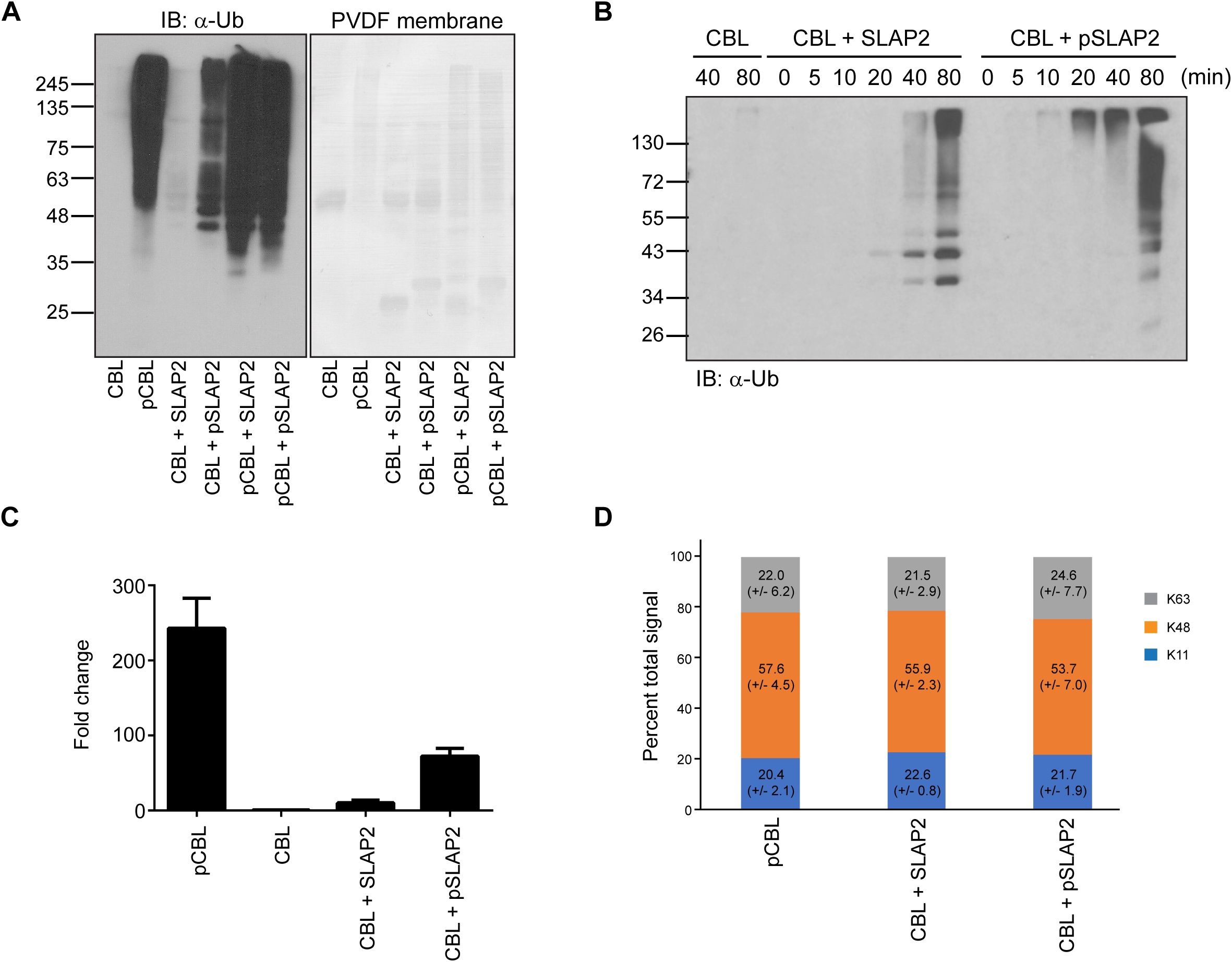
SLAP2 relieves CBL autoinhibition *in vitro*. A) *In vitro* ubiquitination reactions containing CBL TKBD-LHR-RING (CBL) or phosphorylated CBL (pCBL), incubated for 100 min with or without SLAP2 or phosphorylated SLAP2 (pSLAP2), analyzed by SDS-PAGE and immunoblotted with anti-ubiquitin antibody. The right panel shows the Fast Green stained PVDF membrane of the transferred reactions. B) As in A, with reactions stopped over a time course of 80 minutes. C) *In vitro* ubiquitination reactions containing CBL TKBD-LHR-RING (CBL) or phosphorylated CBL (pCBL), incubated for 100 min with or without SLAP2 or phosphorylated SLAP2 (pSLAP2). Histogram showing fold change in E3 ligase activity, measured as polyubiquitin chain production, compared to unphosphorylated CBL using E3LITE assay. The pCBL reaction was diluted 2.7 fold to give a comparable detection range on the plate reader. A representative experiment performed in triplicate is shown with error bars representing standard deviation (SD). Biological replicates with independent protein preparations yielded consistent changes in ubiquitination activity and are shown in Supplementary Fig. 3A. D) *In vitro* ubiquitination reactions were analyzed by LC-MS/MS. Stacked histogram of GG-modified ubiquitin K11, K48, and K63 tryptic peptides, shown as a percentage of total signal intensity of the three peptides detected in each reaction. Values were obtained by peak integration of MS1 ion intensity for each peptide. Shown are average values and standard deviations over three independent experiments.

Ubiquitin contains seven internal lysine residues (K6, K11, K27, K29, K33, K48 and K63) that can be modified with additional Ub molecules to form polyubiquitin chain linkages that determine distinct biological outcomes (Hong, Ng et al., 2015, Kerscher et al., 2006). To further characterize the impact of pSLAP2 on CBL ubiquitination activity, we determined the type of Ub linkages generated *in vitro* by pCBL or CBL in the presence of SLAP2 or pSLAP2. Reactions from the *in vitro* ubiquitination assay were analyzed by LC-MS/MS. Mass spectrometry data was searched against the human proteome, with diglycine as a variable modification on lysine, and the MS1 peak area corresponding to each linkage type quantified (Hong et al., 2015). Activated, phosphorylated CBL generated predominantly K48 linked Ub chains with smaller proportions of K11 and K63 linkages (Fig. 5D). Similar proportions of Ub chain linkages were observed when unphosphorylated CBL was incubated with either SLAP2 or pSLAP2 (Fig. 5D) indicating that the *in vitro* ubiquitin ligase activity of CBL in the presence of SLAP2 or pSLAP2 is qualitatively similar to that of phosphorylated CBL.

To assess the specific role of SLAP2 α-helix in CBL activation, we tested the ability of SLAP2 mutants L241R and S244E to promote CBL ligase activity. Phosphorylated SLAP2 L241R exhibited reduced stimulation of CBL ubiquitination activity compared to pSLAP2 WT by anti-Ub immunoblot (Fig. 6A), and reduced production of polyubiquitin chains in E3 ligase activity assays (Fig. 6B and Supplementary Fig. 3B). Phosphorylated SLAP2 S244E, which maintained CBL binding under low salt conditions (Fig. 2D) such as those used in *in vitro* assays, also exhibited less stimulation of CBL ubiquitination activity than pSLAP2 WT by immunoblot analysis (Fig. 6A) and in the E3 ligase assay (Fig. 6B and Supplementary Fig. 3B). In contrast, SLAP2 mutants R242A, I249W, and S246/250E, which do not interfere with CBL binding but are predicted to disrupt the dimer interface of SLAP2 α-helices observed in our structure, did not interfere with SLAP2 or pSLAP2 activation of CBL (Supplementary Fig. 3C, D). These data confirm that activation of CBL by SLAP2 requires binding of the SLAP2 C-terminal tail α-helix to the CBL TKBD surface identified in the CBL/SLAP2 crystal structure. Given this CBL interface is also involved in maintaining CBL in an inactive state, through interactions with its LHR and RING domain, we refer to this region as the CBL regulatory cleft.

**Figure 6:**
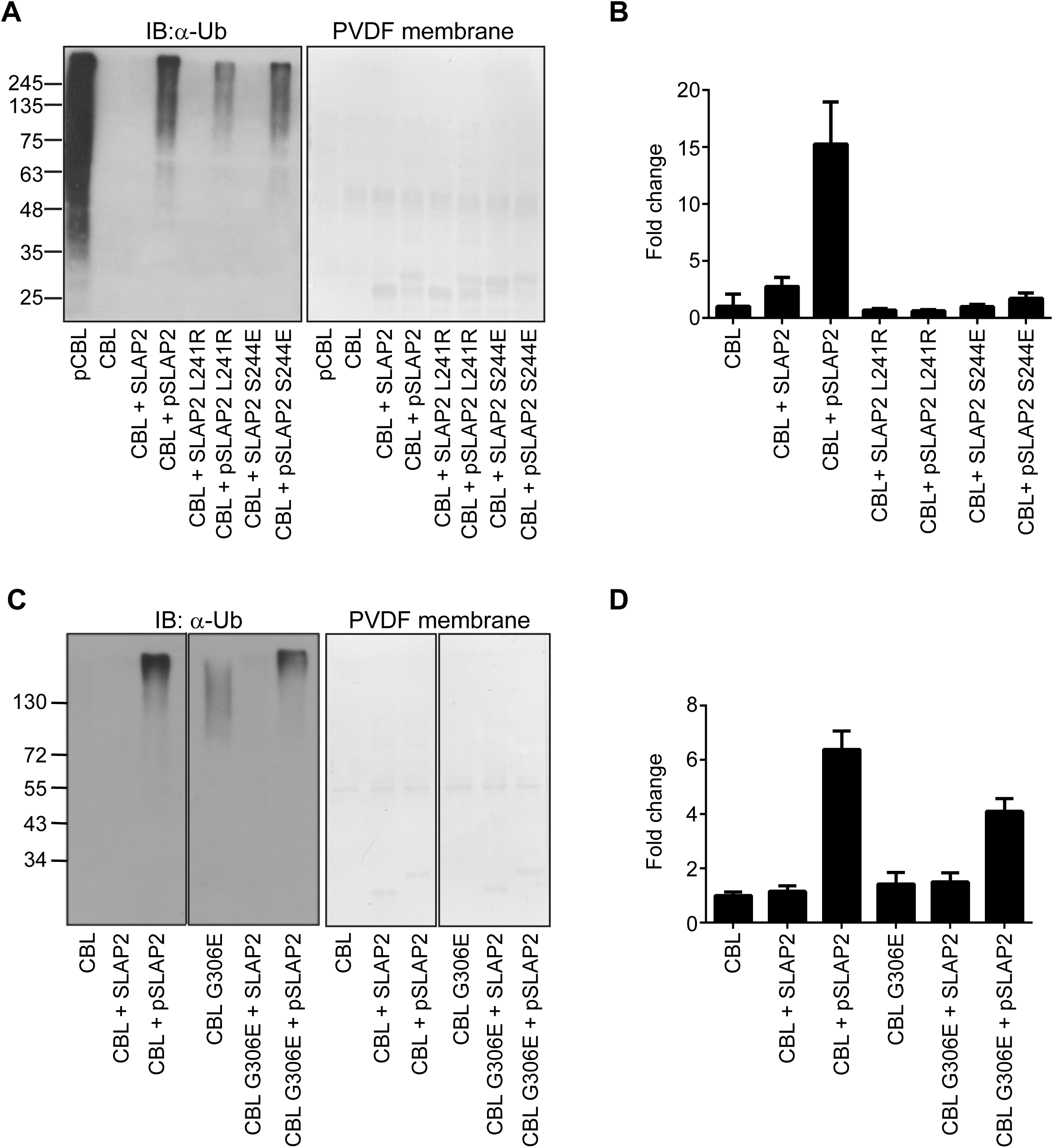
CBL/SLAP2 interface regulates CBL ubiquitination activity. A) *In vitro* ubiquitination reactions containing CBL or pCBL, incubated for 100 min with or without SLAP2 or pSLAP2, WT or mutants (L241R, S244E), analyzed by SDS-PAGE and immunoblotted with anti-Ub antibody. The right panel shows the Fast Green stained PVDF membrane of the transferred reactions. B) Histogram showing fold change in E3 ligase activity in reactions containing CBL and SLAP2 or pSLAP2, WT or mutants, compared to unphosphorylated CBL using E3LITE assay. A representative experiment performed in triplicate is shown with error bars representing SD. Biological replicates with independent protein preparations yielded consistent changes in ubiquitination activity and are shown in Supplementary Fig. 3B. C) and D) As in A and B, respectively, comparing E3 ligase activity of CBL WT and *G306E* incubated with SLAP2 or pSLAP2. Biological replicates with independent protein preparations yielded consistent changes in ubiquitination activity and are shown in Supplementary Fig. 3E.

Engagement of tyrosine phosphorylated substrates by CBL TKBD leads to reorientation of the RING domain and consequently increased E2 binding (Dou et al., 2012). Thus, we tested whether the ability of SLAP2 to promote CBL ubiquitination activity is dependent on canonical TKBD-phospho-tyrosine interactions. CBL mutant *G306E*, which abolishes phospho-tyrosine binding *in vitro* (Meng et al., 1999), behaved similarly to WT CBL, such that pSLAP2 stimulated CBL *G306E* ubiquitination activity (Fig. 6C,D and Supplementary Fig. 3E). This indicates that SLAP2 activation of CBL is independent of both CBL Tyr371 phosphorylation and an intact CBL TKBD substrate binding domain.

### Disruption of the SLAP2 binding TKBD regulatory cleft promotes CBL substrate ubiquitination

Next, the ubiquitination activity of CBL proteins mutated in the CBL regulatory cleft was compared to CBL WT. Strikingly, mutations *A223R, S226E*, and *A223R/S226E* (*AR/SE*) lead to activation of unphosphorylated CBL compared to WT, suggesting that mutations in the SLAP2 binding cleft also disrupt the interaction between CBL TKBD and the LHR (Fig. 7A,B and Supplementary Fig. 3F). These data indicate that mutations of CBL which disrupt SLAP2 binding may also disrupt the autoinhibitory LHR-TKBD interaction and confer activation independent of Y371 phosphorylation.

**Figure 7:**
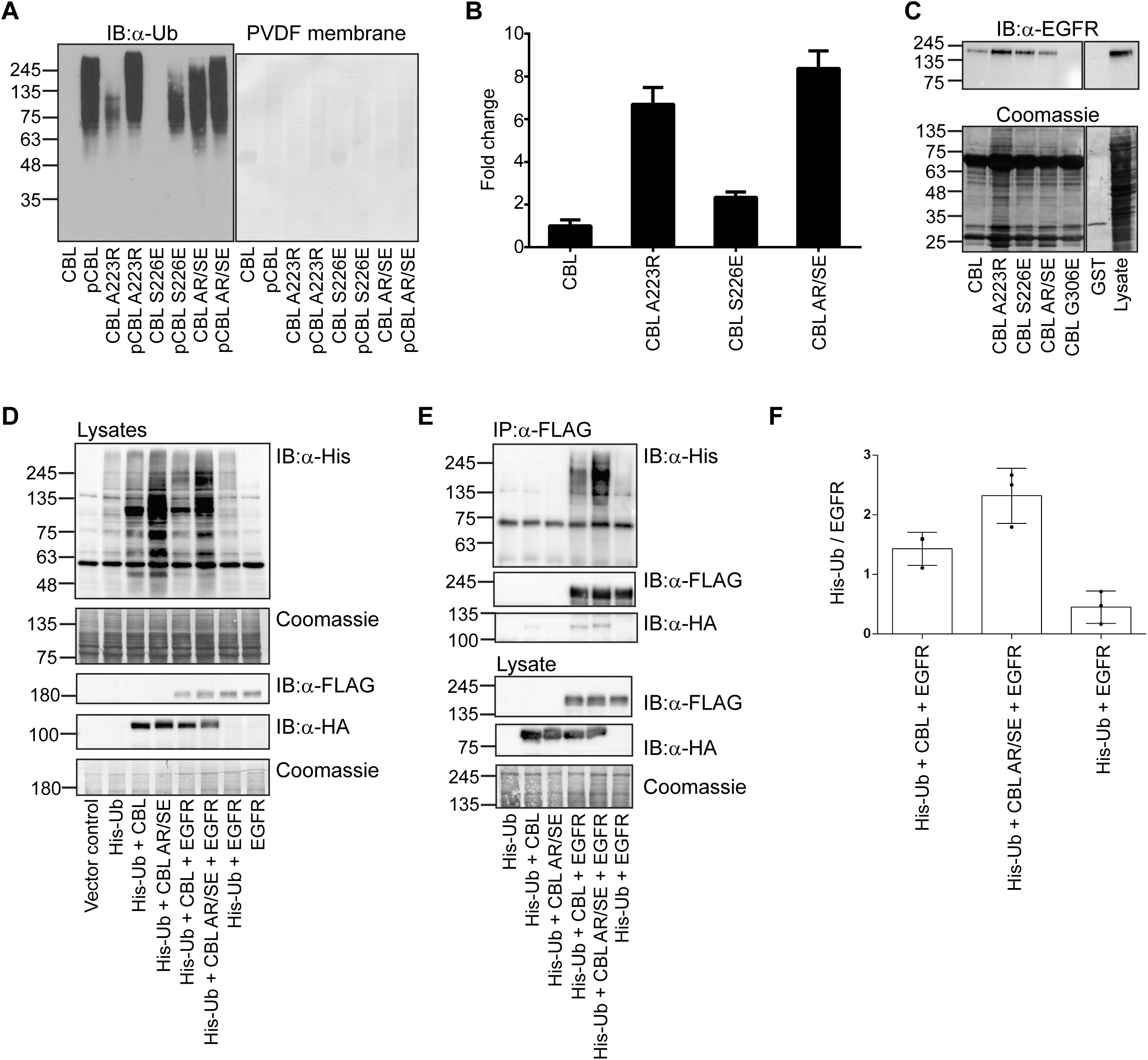
CBL substrate ubiquitination is regulated by the CBL regulatory cleft. A) *In vitro* ubiquitination reactions containing CBL or pCBL WT or mutants (*A223R, S226E*, and double mutant *AR/SE*), analyzed by SDS-PAGE and immunoblotted with anti-Ub antibody. The right panel shows the Fast Green stained PVDF membrane of the transferred reactions. B) Histogram showing fold change in E3 ligase activity of CBL mutants compared to unphosphorylated CBL using E3LITE assay. Error bars represent SD. Biological replicates with independent protein preparations yielded consistent changes in ubiquitination activity and are shown in Supplementary Fig. 3F. C) Immobilized GST-CBL or GST-CBL with TKBD mutations *A223R, S226E* or *AR/SE*, were incubated with EGFR transfected COS-7 cell protein lysates. Bound proteins were resolved by SDS-PAGE and immunoblotted with anti-EGFR. D) Full length HA-CBL WT or *AR/SE* mutant was co-transfected with FLAG-EGFR and His-Ub in HEK 293T cells. Equivalent protein lysates were resolved by SDS-PAGE and immunoblotted with anti-His to detect ubiquitinated species. Coomassie stain of protein loading is shown. Lysates were probed with anti-FLAG and anti-HA to detect FLAG-EGFR and HA-CBL expression. A representative experiment of biological replicates (n=4) is shown. E) Full length HA-CBL WT or *AR/SE* mutant was co-transfected with FLAG-EGFR and His-Ub in HEK 293T cells. EGFR was immunoprecipitated with anti-FLAG, resolved by SDS-PAGE and immunoblotted with anti-His to detect ubiquitinated proteins, anti-FLAG and anti-HA. Expression of transfected EGFR and CBL was detected by anti-FLAG and anti-HA respectively (lower panels). A representative experiment is shown and quantification of biological replicates (n=3) is shown in histogram (F).

The consequences of the CBL *A223R/S226E* regulatory cleft mutations on substrate binding and ubiquitination was examined. First, to confirm that the *AR/SE* mutation does not disrupt TKBD folding and/or binding to phospho-tyrosine substrates, glutathione-S-transferase (GST)-tagged WT or mutant CBL TKBD-LHR-RING fusion proteins were isolated on beads and incubated with lysates from COS-7 cells stimulated with epidermal growth factor (EGF). Captured proteins bound to either CBL WT or regulatory cleft mutants were analyzed by anti-EGF receptor (EGFR) immunoblot (Fig. 7C). CBL mutants *A223R, S226E* and *AR/SE* all retained the ability to bind activated EGFR, indicative of functional TKBD phospho-tyrosine binding, whereas the SH2 mutant *G306E* did not bind EGFR. Next, the *A223R/S226E* mutation was introduced into full length HA-tagged CBL and co-transfected into HEK293 cells with His-Ub and FLAG-tagged EGFR. Protein lysates from cells expressing CBL *AR/SE* showed enhanced incorporation of His-Ub compared to cells expressing CBL WT (Fig. 7D) with or without EGFR co-expression. Furthermore, ubiquitination of immunoprecipitated EGFR was increased when co-expressed with CBL *AR/SE* compared to CBL WT (Fig. 7E, F). Together these data support a role for SLAP2 binding and the TKBD regulatory cleft in promoting CBL ubiquitination of TK substrates.

## Discussion

Compared to other phospho-tyrosine binding domains such as PTB and SH2 domains, the complex architecture of CBL TKBD was suggested to harbour additional protein interaction sites and regulatory regions (Meng et al., 1999). Indeed, the CBL/SLAP2 structure reveals the presence of an additional intermolecular protein interaction surface formed by regions of the SH2 domain, EF-hand and four-helix bundle that is distinct from the phospho-tyrosine binding site. While other adaptor proteins shown to bind CBL TKBD, such as APS and Sprouty, are phospho-tyrosine and SH2 domain dependent, SLAP2 binding involves a unique surface of TKBD and does not require SLAP2 tyrosine phosphorylation (Fong, Leong et al., 2003, Hu & Hubbard, 2005, Wong, Lim et al., 2001). Of note, the SLAP2 binding cleft described in this study overlaps with the TKBD region that was previously described as part of a conserved binding channel and predicted to provide an additional protein interaction region (Zheng et al., 2000).

CBL E3 ligase function is integral to regulation of tyrosine kinase signaling through ubiquitination of tyrosine phosphorylated substrates. Furthermore, CBL activity is controlled by tyrosine phosphorylation which regulates the equilibrium between the autoinhibited and catalytically open conformational states (Dou et al., 2012). Our study reveals that SLAP/SLAP2 adaptor binding provides an additional mechanism to regulate CBL E3 ligase activity. The structure of SLAP2 bound to CBL TKBD showed that the SLAP2 binding site overlaps the region bound by the LHR in the autoinhibited conformation (Dou et al., 2012). Therefore, SLAP2 adaptor binding could hinder access to the closed conformation, shifting the equilibrium toward the activated state. Indeed, addition of SLAP2 promoted CBL ubiquitin ligase activity *in vitro*. However, SLAP2 binding did not stimulate CBL activity to the same magnitude as phosphorylation of CBL at Tyr371. This suggests that the effect of SLAP2 binding on the equilibrium between closed and open conformations is dynamic, presumably due to the absence of phosphorylated LHR residue Tyr371, which serves to stabilize the active conformation of CBL (Dou et al., 2012). While the aromatic ring of pTyr371 forms hydrophobic interactions with other LHR residues in activated CBL, the phosphate moiety forms crucial hydrophilic interactions with Lys382 and Lys389 of the RING domain (Dou et al., 2012). Therefore, SLAP2 interaction with CBL would favor an open, catalytically competent conformation, but lack the pTyr371 interactions with the RING domain that further stabilize E2 bound ubiquitin for transfer (Dou et al., 2012, Dou et al., 2013).

It remains unclear why phosphorylation of SLAP2 Y248, although not required for binding to TKBD, lead to more robust CBL activation *in vitro*. Since Y248 resides within the region that binds the TKBD, phosphorylation at Y248 may stabilize the SLAP2 α-helix or facilitate order transition within the SLAP2 carboxy-terminal region, as observed for numerous signaling proteins (Dunker, Oldfield et al., 2008, Iakoucheva, Radivojac et al., 2004). Phosphorylation at Y248 may also increase the affinity of SLAP2 for CBL, thus displacing the LHR from the autoinhibited conformation more efficiently. Our assessment of binding affinity through standard approaches was hindered due to the hydrophobic and linear nature of corresponding synthetic peptides and the instability and insolubility of purified SLAP2 proteins at concentrations and conditions required for quantitative binding assays. As well, it is possible that phosphorylation at Y248 results in allosteric regulation of CBL activity or impacts how full-length SLAP2 protein interacts with CBL. Additional investigation will be needed to address the dynamic conformational changes in CBL TKBD-LHR-RING induced by SLAP2 and pSLAP2 proteins.

Three mammalian CBL proteins are encoded by separate genes: CBL (also known as c-Cbl and studied here), Cbl-b, and Cbl-c (Nau & Lipkowitz, 2003). CBL and CBL-b proteins are highly homologous in the TKBD and Cbl-b is also regulated by phosphorylation of LHR tyrosine residue (Y363), which relieves autoinhibition to enhance ubiquitin ligase activity (Dou et al., 2012, Dou et al., 2013, Keane, Rivero-Lezcano et al., 1995, Kobashigawa, Tomitaka et al., 2011). The residues involved in SLAP2 binding as identified by our CBL/SLAP2 crystal structure are conserved in Cbl-b TKBD and SLAP has been shown to bind Cbl-b *in vitro* (Sharma, Ponce et al., 2019). Therefore, SLAP binding may also provide a mechanism for phosphorylation independent activation of Cbl-b.

Protein-protein interactions as a means to regulate activity has previously been observed for E3 ligases, including CBL family proteins. Cbl-c ubiquitination activity is increased by interaction of its RING finger domain with a LIM domain of Hic-5, a member of the paxillin family(Ryan, Kales et al., 2012). However, in contrast to our observations for activation of CBL by SLAP2 binding, Hic-5 can increase the activity of Cbl-c only after the E3 is activated by phosphorylation or phosphomimetic mutation at LHR Y341(Ryan et al., 2012). The HECT E3 Smurf2 also adopts an autoinhibited structure that is relieved by binding an adaptor protein, Smad7(Ogunjimi, Briant et al., 2005, Wiesner, Ogunjimi et al., 2007). Smad7 regulates the ubiquitin ligase activity of Smurf2 by promoting E2 binding to the HECT domain (Ogunjimi et al., 2005). Thus, adaptor protein binding as a means to regulate E3 activity may be a more general mechanism (Buetow & Huang, 2016, Kales, Ryan et al., 2012).

Our data help explain the requirement for SLAP/SLAP2 adaptor proteins in CBL dependent downregulation of TK signaling. Analysis of SLAP deficient mice have highlighted its involvement in T and B cell development, dendritic cell maturation, platelet activation and mast cell degranulation (Cherpokova, Bender et al., 2015, Dragone, Myers et al., 2006b, Liontos et al., 2011, Sharma et al., 2019, Sosinowski et al., 2001). The phenotypes observed were shown to be a consequence of defective down regulation of tyrosine kinase linked receptors including TCR, BCR, GM-CSFR (Dragone et al., 2006b, Liontos et al., 2011, Sosinowski et al., 2001). In addition, studies in cell lines have demonstrated that the SLAP/SLAP2 C-terminus is required for CBL dependent downregulation of TCR and BCR as well as receptor tyrosine kinases such as CSF-1R and FLT3 (Dragone et al., 2006a, Holland et al., 2001, Kazi & Ronnstrand, 2012, Loreto et al., 2002, Myers, Dragone et al., 2005, Myers et al., 2006, Pakuts et al., 2007, Wybenga-Groot & McGlade, 2015). These studies have suggested that SLAP/SLAP2 adaptor function has a role in the recruitment of CBL to activated tyrosine kinase complexes, facilitating phosphorylation and proximity to substrates. Our results reveal a more complex role in which the SLAP/SLAP2 C-terminal region binding to CBL TKBD also regulates ubiquitin ligase activity. Developmentally regulated and cell type specific expression of SLAP/SLAP2, therefore, could promote CBL activity in specific cellular contexts (Dragone et al., 2009, Lebigot et al., 2003, Wybenga-Groot & McGlade, 2015).

SLAP/SLAP2 binding to CBL may also allow the ubiquitination of substrates in the absence of CBL activation mediated by tyrosine phosphorylation. Such a mechanism could contribute to the constitutive ubiquitination of TCR CD3ξ chain in double positive (DP) thymocytes that occurs during T-cell development. A prior study demonstrated that MHC-independent tonic ubiquitination of the TCR:CD3 complex requires both CBL and SLAP and coincides with the timing of upregulated SLAP expression in DP thymocytes (Sosinowski et al., 2001, Wang, Holst et al., 2010). In this context, SLAP binding may serve to promote ubiquitination of substrates in the absence of activated TK signaling and CBL tyrosine phosphorylation by favoring the catalytically competent conformation. Alternatively, in the presence of an activated tyrosine kinase, SLAP/SLAP2 binding and displacement of the TKBD bound LHR might facilitate Y371 phosphorylation and lower the threshold for CBL activation.

In addition to the role of the SLAP/SLAP2 carboxy-terminal tail in CBL ubiquitin ligase activity, binding induced conformational changes in CBL may also influence substrate selection, as well as protein-protein interactions related to CBL adaptor function. Although we did not observe any differences in ubiquitin chain linkages assembled by phosphorylated CBL or SLAP bound CBL *in vitro*, it is possible that distinct mechanisms of activation could influence substrate ubiquitination qualitatively, resulting in distinct protein fates. In addition, the open conformation induced by SLAP/SLAP2 binding may be similar to that imposed by some LHR mutations which allow RING flexibility and ubiquitination of substrates bound to regions distinct from TKBD phospho-tyrosine substrate binding pocket (Buetow, Tria et al., 2016). Together, our observations suggest that SLAP/SLAP2 binding to CBL could provide a mechanism to tune CBL activity or substrate selection in the context of different degrees of tyrosine kinase signaling.

Mutations in *CBL* which impair E3 ligase activity have been identified in juvenile myelomonocytic leukemia (JMML), chronic myelomonocytic leukemia (CMML), and chronic myeloid leukemia (CML), as well as in patients with several other myeloproliferative neoplasms (Grand, Hidalgo-Curtis et al., 2009, Kales et al., 2010, Loh et al., 2009, Makishima, Cazzolli et al., 2009, Ogawa et al., 2010, Swaminathan & Tsygankov, 2006, Thien & Langdon, 2005). In addition to mutations in the RING domain, the LHR, and Tyr371 in particular, are frequent targets of missense mutations that prevent phosphorylation dependent activation of CBL ubiquitin ligase function (Dou et al., 2012, Kales et al., 2010, Loh et al., 2009, Ogawa et al., 2010). These mutant forms of CBL are also thought to act through a gain-of-function mechanism, since adaptor functions are retained, allowing signaling protein recruitment through multiple carboxy-terminal SH2 and SH3 binding motifs (Javadi, Richmond et al., 2013, Nadeau, An et al., 2017). Understanding of this alternative activation mechanism which bypasses Y317 phosphorylation provides insights that may be used to target CBL activation and restore E3 ligase activity.

In conclusion, our results reveal a novel phosphorylation independent protein-protein interaction interface on the CBL TKBD and demonstrate that SLAP/SLAP2 adaptor protein binding to this site is an additional mechanism to regulate CBL ubiquitin ligase activity. Furthermore, the presence of SH2 and SH3 domains as well as a myristoylation modification in SLAP/SLAP2 support a model in which SLAP/SLAP2 binding likely regulates additional context specific aspects of CBL function.

## Methods

### Protein overexpression and purification

Mouse SLAP2 (residues 28-259) and CBL TKBD (residues 25-357) proteins were overexpressed and purified as a complex as described elsewhere (Wybenga-Groot & McGlade, 2020; https://biorxiv.org/cgi/content/short/2020.10.08.331397v1). Residues 29-261 of human SLAP2 or 19-254 of mouse SLAP were cloned in frame into a modified pET32a vector (Wybenga-Groot & McGlade, 2020) to express thioredoxin-tagged (Trx)-His(6)-hSLAP2, and - mSLAP. Using standard QuikChange methods, 19 residues downstream of the TEV cleavage site were deleted from Trx-His(6)-hSLAP2, thus abolishing the thrombin cleavage site and placing the TEV cleavage site in closer proximity to the protein N-terminus, to generate Trx-His(6)-Δlinker-hSLAP2. Trx-His(6)-Δlinker-hSLAP2 or Trx-His(6)-mSLAP were cloned in frame into multiple cloning site (MCS) 1 of pETDuet-1 (Novagen), with CBL (47-357) cloned in frame into MCS2 of the same pETDuet-1 vector, for co-expression of SLAP2 or SLAP and CBL (Duet-His-Δlinker-hSLAP2-CBL and Duet-His-mSLAP-CBL). Residues 2-436 of CBL were cloned in frame into pGEX4T-1 to express glutathione-S-transferase (GST)-CBL TKBD-LHR-RING. GST-EphA4 (591-896) plasmid was a gift from Frank Sicheri (Lunenfeld-Tanenbaum Research Institute)(Wybenga-Groot, Baskin et al., 2001). Mutations of residues were generated by standard QuikChange site-directed mutagenesis methods.

For co-expression of CBL and SLAP2/SLAP, Duet-His-Δlinker-hSLAP2-CBL WT, Duet-His-mSLAP-CBL WT, and their respective point mutants were overexpressed in *Escherichia coli* BL-21 or TKB1 competent cells (Stratagene) (100 mL Luria Bertani (LB) media supplemented with 50 µg/ml ampicillin, overnight at 16-18°C, and 0.35-0.5 mM IPTG induction, with modifications for TKB1 cells as per manufacturer’s instructions). Cell pellets were resuspended in either high salt (50 mM HEPES pH 7.5, 0.5 M NaCl, 10% glycerol, 10 mM imidazole, 10 mM β-mercaptoethanol (βME), 10 mM MgCl_2_, 5 mM CaCl_2_, cOmplete EDTA-free protease inhibitor cocktail tablets (inhibitor tablets) (Roche Applied Science), benzonase nuclease, 1 mM phenylmethylsulfonyl fluoride (PMSF)) or low salt (as for high salt except 25 mM HEPES pH 7.5, 150 mM NaCl) lysis buffer, lysed by sonification, clarified by centrifugation, mixed with Ni-NTA agarose (Qiagen), and resin washed with 25 mM HEPES pH 7.5, 150 mM NaCl, 2% glycerol, 20 mM imidazole, 10 mM βME. Bead slurry was mixed with SDS 2X loading buffer, boiled, and analyzed by 10 or 12.5% SDS-PAGE, stained with Coomassie blue.

Purification of Trx-His(6)-mSLAP2 for *in vitro* phosphorylation experiments was performed as described elsewhere (Wybenga-Groot & McGlade, 2020). GST-EphA4 was overexpressed (2L LB, overnight at rt, 0.15 mM IPTG induction) and cell pellets resuspended in lysis buffer (50 mM HEPES pH 7.5, 0.5 M NaCl, 10% glycerol, 3 mM DTT, 1 mM MgSO_4_, inhibitor tablets, benzonase nuclease). Following sonification and centrifugation, supernatant was mixed with Glutathione Sepharose 4B (GE Healthcare) for 90 min by nutating at 4°C. Resin was washed with 4 x 25 mL of PBS, pH 7.4, 3 mM DTT.

For purification of SLAP2, pSLAP2, CBL-TKBD-LHR-RING (CBL) and pCBL for ubiquitination assays, GST-CBL-TKBD-LHR-RING and Trx-His(6)-hSLAP2 WT and point mutants were transformed into *E. coli* BL-21 or TKB1 as above. 100 mL cultures were grown overnight at 18°C (A_600_ = 0.6-0.9 and 0.4 mM IPTG induction), pelleted and stored at -80°C. CBL pellets were processed as above using NP-40 lysis buffer (50 mM HEPES pH 7.5, 150 mM NaCl, 10% glycerol, 1% NP-40, 10 mM NaF, 1.5 mM MgCl_2_, 1 mM EDTA pH 8.0, 1 mM Na_3_VO_4_, and inhibitor tablets) and NP-40 wash buffer (20 mM HEPES pH 7.5, 150 mM NaCl, 5% glycerol, 0.1% NP-40, 5 mM βME, 0.5 PMSF, 0.5 mM Na_3_VO_4_). After three washes with PBS wash buffer (PBS pH 7.4, 5 mM βME, 0.5 mM Na_3_VO_4_), resin was incubated with thrombin protease (Sigma #T4648) and CBL protein eluted with PBS. SLAP2 pellets were resuspended in 3 mL each of lysis buffer (50 mM HEPES pH 7.5, 0.5 M NaCl, 10% glycerol, 1% NP-40, 10 mM imidazole, 10 mM NaF, 5 mM βME, 5 mM caproic acid, 1.5 mM MgCl_2_, 1 mM Na_3_VO_4_, benzonase nuclease, and inhibitor tablets) and processed as above. SLAP2 and pSLAP2 were isolated using Ni-NTA agarose resin followed by three washes with 50 mM HEPES pH 7.5, 0.5 M NaCl, 2% glycerol, 20 mM imidazole, 5 mM βME, 0.5 mM Na_3_VO_4_, three washes with PBS wash buffer, and incubation with TEV protease. Protein concentrations were determined through Bradford assays.

### Structure determination

The C-terminal tail of mouse SLAP2 (residues 237-255) and CBL TKBD was crystallized as a complex and details are described elsewhere (Wybenga-Groot & McGlade, 2020). The structure of the complex was determined by molecular replacement using CBL TKBD (PDB id: 1B47) as a model, followed by iterative cycles of model building in Coot and Phenix, and refinement with Phenix_refine (Adams, Afonine et al., 2010, Emsley & Cowtan, 2004, Wybenga-Groot & McGlade, 2020). Atom-atom distances, superpositions, and root mean square deviations were calculated in Coot (Emsley & Cowtan, 2004). Accessible surface area was calculated with Areaimol from the CCP4 program suite (Winn, Ballard et al., 2011). Coordinates and structure factors were deposited in the Protein Data Bank with accession number 6XAR.

### *In vitro* phosphorylation and detection by mass spectrometry

Purified mSLAP2 was incubated with GST-EphA4, 50 mM Hepes pH 7.5, 20 mM MgCl_2_, 5 mM adenosine triphosphate (ATP), 5 mM DTT, and 1 mM Na_3_VO_3_ overnight at rt. Reaction was boiled with SDS 2X loading buffer, loaded on 12.5% SDS-PAGE gel, and transferred to polyvinylidene difluoride (PVDF) membrane. PVDF membrane was immunoblotted with anti-pTyr antibody 4G10 (Upstate Biotechnology, Inc.) as per manufacturer’s protocol. A second kinase reaction was boiled with SDS 2X loading buffer and run in multiple lanes of a 15% SDS-PAGE gel. The gel was stained with Coomassie and the upper band observed in the presence of EphA4 excised from the gel. Gel bands were treated and digested with either trypsin or GluC protease as per SPARC BioCentre’s in-gel digestion protocol (https://lab.research.sickkids.ca/sparc-molecular-analysis/services/mass-spectrometry/mass-spectrometry-sample-protocols/). Digested peptides were subjected to LC-MS/MS at SPARC BioCentre (The Hospital for Sick Children) (60 min gradient, Thermo LTQ Orbitrap) and the raw data searched with PEAKS software against the mouse proteome, with carbamidomethylation as a fixed modification and deamidation (NQ), oxidation (M), and phosphorylation (STY) as variable modifications.

### *In vitro* autoubiquitination assays

*In vitro* ubiquitination reactions were prepared with 20 pmol of CBL or pCBL and 60 pmol of hSLAP2 or phosphorylated hSLAP2 (pSLAP2) for both WT and mutants. Reagents were thawed and reactions prepared on ice. CBL and SLAP2 were mixed in PBS (with 5 mM βME, 5 mM caproic acid, 0.5 mM PMSF, and 0.5 mM Na_3_VO_4_) at rt for 5-10 min before addition of master mix (42 pmol E1 (UBE1, Boston Biochem or Ubiquitin-Proteasome Biotechnologies), 0.1 μmol E2 (UbcH5b/UBE2D2, Boston Biochem #E2-622), 1 mmol ATP, and 11.6 μmol ubiquitin (Boston Biochem #U-100H) in reaction buffer (0.2 M Tris pH 7.5, 10 mM MgCl_2_, 2 mM DTT). Reactions were incubated at 30°C shaking at 500-700 rpm for 60-110 minutes and terminated by addition of SDS 2X loading buffer and boiling for 5-10 minutes. Equal volumes of each reaction were loaded on 12.5% SDS-PAGE gel, transferred to PVDF membrane, and immunoblotted for ubiquitin as per standard protocols (anti-ubiquitin antibody P4G7, BioLegend #838701, 1:1000; anti-mouse IgG HRP-linked antibody, Cell Signaling Technology #7076, 1:10000; Western Lightning Plus ECL, Perkin Elmer).

Ubiquitination reactions were set up as above (25 µL total volume) in triplicate and diluted with reaction buffer after completion (1:40 for pCBL, 1:15 for all other reactions). Each reaction triplicate was added in triplicate to individual wells of a 384-well E3LITE Customizable Ubiquitin Ligase Kit (Life Sensors) plate, prewashed three times with PBS, and incubated for 60 minutes at rt on a rotating platform to capture ubiquitinated proteins. Wells were washed three times with PBS plus 0.1% Tween (PBS-T) and 25 uL of detection solution 1 (1:1000 dilution in PBS-T + 5% BSA) added to each well. After 50 min incubation, the wells were washed three times with PBS-T, 25 μL of streptavidin peroxidase polymer ultrasensitive antibody (Sigma #S2438, 1:10000 dilution in PBS-T, 5% BSA) added to each well, and the plate incubated for 50 min. Wells were washed four times with PBS-T and 25 μL of prepared Immobilon Western Chemiluminescent HRP Substrate (Millipore #P90719) added to each well immediately prior to the plate being read on a Synergy Neo2 Plate Reader (BioTek Instruments). The average luminescent signal for each condition was divided by that of CBL WT to determine fold change over CBL. Standard deviation of the reaction averages was calculated and used as a measure of error. Each reaction set was analyzed at least twice with fresh protein purifications.

### LC-MS analysis of ubiquitination reactions

Relative ubiquitin linkages levels were measured by LC-MS via monitoring of diglycine(GG)-modified ubiquitin “linkage reporter” peptides (Hong et al., 2015). *In vitro* autoubiquitination reactions were prepared as above, lyophilized, re-suspended in 9 M urea, 100 mM NH_4_HCO_3_, 5 mM DTT, and incubated for 20 min at 60°C. Samples were cooled to rt, treated with 10 mM iodoacetamide for 30 min at rt in the dark, and diluted with 50 mM NH_4_HCO_3_ to a final urea concentration of ∼2M. Sequencing-grade, TPCK-treated, modified trypsin (Promega) was added (1 µg) for 16 hrs at 37°C. The resulting peptide samples were desalted using C18 tips and lyophilized. Peptides were re-suspended in 0.1% HCOOH and analyzed by LC-MS/MS. LC was conducted using a C18 pre-column (Acclaim PepMap 100, 2 cm x 75 µm ID, Thermo Scientific) and a C18 analytical column (Acclaim PepMap RSLC, 50 cm x 75 µm ID, Thermo Scientific) over a 120 min reversed-phase gradient (0-40% ACN in 0.1% HCOOH) at 225 nl/min on an EASY-nLC1200 pump (Proxeon) in-line with a Q-Exactive HF mass spectrometer (Thermo Scientific). A MS scan was performed with a resolution of 60,000 followed by up to 20 MS/MS scans (minimum ion count of 1000 for activation) using higher energy collision induced dissociation (HCD) fragmentation. Dynamic exclusion was set for 5 seconds (10 ppm; exclusion list size = 500). For peptide and protein identification, Thermo.RAW files were converted to.mzML format using ProteoWizard (v3.0.10800; (Kessner, Chambers et al., 2008)), then searched using X!Tandem (X!TANDEM Jackhammer TPP v2013.06.15.1; (Craig & Beavis, 2004)) and Comet (v2014.02 rev.2; (Eng, Jahan et al., 2013)) against the human RefSeq v45 database (containing 36113 entries). Search parameters specified a parent ion mass tolerance of 10 ppm and an MS/MS fragment ion tolerance of 0.4 Da, with up to two missed cleavages allowed for trypsin (excluding K/RP). Variable modifications of 0.984013 on N/Q, 15.99491 on M, 114.04293 on K, 79.96633 Y/S/T and 57.021459 on C were allowed. Data were filtered through the TPP (v4.7 POLAR VORTEX rev 1) with general parameters set as –p0.05 -x20 –PPM. For label-free MS1-level quantification, raw files were analyzed using MaxQuant (v1.6; (Cox & Mann, 2008)).

### Ubiquitination and substrate binding in cells

COS-7 cells were serum starved for 24 hours (∼80% confluent), followed by a 15 minute stimulation with 100 ng/mL EGF. Cells were lysed in 200 µL PLC lysis buffer/plate (50 mM HEPES pH 7.5, 150 mM NaCl, 1.5 mM MgCl_2_, 1 mM EGTA pH 8.0, 10% glycerol, 1% Triton X-100, supplemented with 10 mM NaF, Na_3_VO_4_, cOmplete EDTA-protease inhibitor tablet (Roche)) and protein concentration determined by Bradford assay. Lysate (500 µg) was incubated for 2 hours at 4°C with the equivalent of 30 µg GST-CBL-TKBD-LHR-RING WT or mutants captured on Glutathione Sepharose 4B as above. Samples were prepared for SDS-PAGE and immunoblot as described above, with anti-EGFR antibody (D38B1; Cell Signaling Tech). HEK 293T cells were transfected using Lipofectamine 2000 (ThermoFisher), as per manufacturer’s instructions and standard procedures, with 3 µg HA-tagged full length CBL (in pCDEF3), 1 µg FLAG-tagged EGFR (in pcDNA 3.1), and/or 3 µg His-Ubiquitin (octameric ubiquitin construct with N-terminal His_6_ tag for each), as indicated. Promoter matched backbone DNA was used where necessary to give 7 µg of DNA per plate for all conditions. After 24 hours, cells were harvested in cold PBS, pelleted by centrifugation, lysed in cold PLC lysis buffer, and protein concentration measured by Bradford assay. For lysate immunoblots, 50 µg samples were prepared in 2X SDS loading buffer and boiled for 10 min. For immunoprecipitation, 1 mg of lysate was incubated with pre-washed anti-FLAG M2 affinity gel (Sigma-Aldrich, #A2220) for 2 hours at 4°C with gentle nutation. Resin was pelleted at 5000 rcf for 30 seconds at 4°C and washed three times with lysis buffer. After the last wash, resin was mixed with 70 μL of 2X SDS loading buffer and boiled for 10 minutes. For immunoblotting, immunoprecipitation samples were divided between two gels. Lysate and immunoprecipitation samples were resolved by 8% SDS-PAGE and immunoblotted using standard protocols as described above. Where indicated, the following primary antibodies were used: 1 μg/mL anti-FLAG M2 (Sigma-Aldrich #F1804), 0.2 µg/mL anti-HA F-7 (Santa Cruz Biotechnology #sc7392), 0.5 µg/mL anti-His_6_ (Roche #11 922 416 001). Total protein on lysate membranes was stained with 50% methanol with 0.1% Coomassie Brilliant Blue R-250 dye, and destained in 50% methanol, 10% acetic acid. The His-Ub, FLAG (EGFR) signals were quantified for the FLAG immunoprecipitation using Image Lab (BioRad Laboritories, version 6.0 build 26). Lanes and bands were manually defined using the volume tool. Background adjusted volume measurements were obtained for the FLAG and His signals for three replicates of the FLAG immunoprecipitation (IP) experiment in conditions where FLAG-EGFR was transfected. His-Ub signals were normalized to the corresponding FLAG signal for each replicate. The average normalized intensities were plotted, showing the corresponding individual values as points, and error bars representing the standard deviation.

## Supporting information

Supplementary tables and figures and PDB validation report

## Acknowledgements

The authors thank Savar Kaul, Julia Manalil, Donna Berry, Jessica Lapierre and Andrew Bondoc for technical assistance, and Jean-Francois Cote for comments on the manuscript. This work was supported with funds from the Cancer Research Society, Garron Family Cancer Centre, and the Canadian Institutes of Health Research FRN166034(to CJM). LWG was support by fellowships from CIHR and the American Brain Tumor Association, AT is funded by an Ontario Graduate Scholarship and RESTRACOMP.

## Author Contributions

LEW-G, AT and CS performed experiments and wrote the manuscript; JSG performed experiments; MM supervised and BR supervised JSG. CJM supervised LEW-G, CS, AT and wrote the manuscript.

## Competing Interest Statement

The authors declare no conflict of interest.

## Notes

### Competing Interest Statement

The authors have declared no competing interest.

