## Supplementary tables and figures and PDB validation report for "SLAP2 adaptor binding disrupts c-CBL autoinhibition to activate ubiquitin ligase function"

Supplementary Information.

**Supplementary Table 1 – Distances between CBL and SLAP2 atoms in CBL/SLAP2 crystal structure**

| SLAP2 Molecule 1 | CBL Molecule 1 | CBL Secondary Structure Element | Distance (Å) |
|---|---|---|---|
| Leu 237 O | Val 263 O | αN-βA loop | 2.8 |
| Leu 237 CD1 | Arg 343 NH2 | SH2 C-terminus | 3.4 |
| Leu 237 CB | Thr 264 CA | αN-βA loop | 4.6 |
| Glu 239 OE1 | Ala 262 O | helix αN | 3.0 |
| Glu 239 OE1 | Val 263 O | αN-βA loop | 2.8 |
| Glu 239 OE2 | Ala 262 O | helix αN | 2.8 |
| Gly 240 CA | Met 269 CA | strand βA | 4.4 |
| Gly 240 O | Ala 270 CB | strand βA | 3.8 |
| Leu 241 CD1 | Trp 258 CZ2 | helix αN | 4.0 |
| Leu 241 CD1 | Val 263 CG2 | αN-βA loop | 3.6 |
| Leu 241 CD2 | Ala 223 CB | helix αE2 | 3.7 |
| Leu 241 N | Tyr 268 CE2 | strand βA | 4.1 |
| Ser 244 OG | Ser 226 OG | helix αE2 | 3.8 |
| Leu 245 CG | Ala 223 CA | helix αE2 | 4.3 |
| Leu 245 CD2 | Met 222 CE | helix αE2 | 3.7 |
| Tyr 248 OH | Lys 225 NZ | helix αE2 | 3.2 |
| Tyr 248 CD1 | Cys 232 SG | αE2-αF2 loop | 4.8 |
| Leu 251 CD1 | Lys 153 CB | helix αD | 3.7 |
| Leu 251 CD2 | Gln 128 NE2 | helix αC | 3.8 |
| Leu 251 CD2 | Asn 150 OD1 | helix αD | 3.6 |
| Leu 251 CD2 | Leu 154 CD2 | helix αD | 4.7 |
| Leu 251 O | Gln 128 NE2 | helix αC | 3.2 |
| Ala 252 CB | Cys 232 SG | αE2-αF2 loop | 4.2 |

**Supplementary Table 2 – Distances between SLAP2 atoms in CBL/SLAP2 crystal structure**

| SLAP2 Molecule 1 | SLAP2 Molecule 2 | Distance (Å) |
|---|---|---|
| Arg 242 NE | Ile 249 O | 3.3 |
| Arg 242 NH2 | Ala 252 O | 2.5 |
| Arg 242 NH1 | Ser 250 O | 2.9 |
| Arg 242 NE | Asp 254 OD | 3.1 |
| Leu 245 CB | Ile 249 CG2 | 4.1 |
| Leu 245 CB | Ile 249 CD1 | 3.7 |
| Leu 245 CD1 | Ile 249 CD1 | 3.9 |
| Ser 246 OG | Ser 250 OG | 3.2 |
| Ile 249 CG2 | Leu 245 CB | 3.9 |
| Ile 249 CD1 | Leu 245 CB | 3.7 |
| Ile 249 CD1 | Leu 245 CD1 | 3.8 |
| Ile 249 O | Arg 242 NH1 | 2.6 |
| Ser 250 OG | Ser 246 OG | 4.2 |
| Ala 252 O | Arg 242 NH2 | 3.2 |
| Asp 254 OD2 | Arg 242 NH1 | 2.6 |

**Supplementary Table 3 – Distances between CBL atoms in CBL/SLAP2 crystal structure**

| CBL Molecule 1 | CBL Molecule 2 | Distance (Å) |
|---|---|---|
| Glu 135 CA | Phe 284 CZ | 8.1 |
| Glu 135 CA | Lys 283 CA | 10.2 |
| Lys 137 CA | Lys 283 CA | 11.3 |
| Glu 143 O | Glu 143 O | 3.8 |
| Asn 144 CB* | Ser 145 CA | 4.5 |
| Asn 144 ND2* | Gln 146 N | 3.6 |
| Ser 145 CA | Asn 144 CB | 5.3 |
| Gln 146 N | Asn 144 ND2* | 4.3 |
| Leu 219 CA | Leu 219 CD2 | 3.8 |
| Leu 219 CD2 | Leu 219 CA | 3.8 |
| Leu 219 CD2 | Met 222 SD | 3.6 |
| Met 222 SD | Leu 219 CD2 | 4.4 |
| Met 222 SD | Met 222 SD | 5.9 |
| Lys 283 CA | Lys 137 CA | 11.1 |
| Lys 283 CB | Glu 135 CA | 7.4 |
| Phe 284 CZ | Glu 135 CA | 9.0 |

*Electron density absent for these atoms

**Supplementary Figure 1: CBL/SLAP2 binding interface**

Ribbon representation of the Cα atoms of the CBL/SLAP2 crystal structure, with the 4H bundle, EF-hand, and SH2 domain coloured teal, dark blue, and light blue, respectively. SLAP2 is coloured green. Side chains shown in stick are coloured according to their respective backbones. Oxygen, nitrogen, and sulfur atoms are coloured red, blue, and yellow, respectively.

**Supplementary Figure 2: Magnification of CBL/SLAP2 interface**

Representation of the Cα atoms of the CBL/SLAP2 crystal structure, with CBL molecules coloured light and dark blue and SLAP2 molecules coloured light and dark green, with side chains coloured according to their respective backbones. Oxygen and nitrogen atoms are coloured red and blue, respectively.

**Supplementary Figure 3: Ubiquitination activity *in vitro***

A) Histogram showing fold change in ubiquitination activity of pCBL or CBL plus SLAP2 or pSLAP2 compared to unphosphorylated CBL using E3LITE assay. The pCBL reaction was diluted 2.7 fold to give a comparable detection range on the plate reader. A representative experiment performed in triplicate is shown with error bars representing SD. B) As in A), for SLAP2 WT and mutants. C) *In vitro* ubiquitination reactions containing CBL or pCBL, incubated with or without SLAP2 or pSLAP2, WT or mutants, analyzed by SDS-PAGE and immunoblotted with anti-Ub antibody. D-F) As in A), for CBL and SLAP2 WT and mutants.

**Supplementary Figure 1**

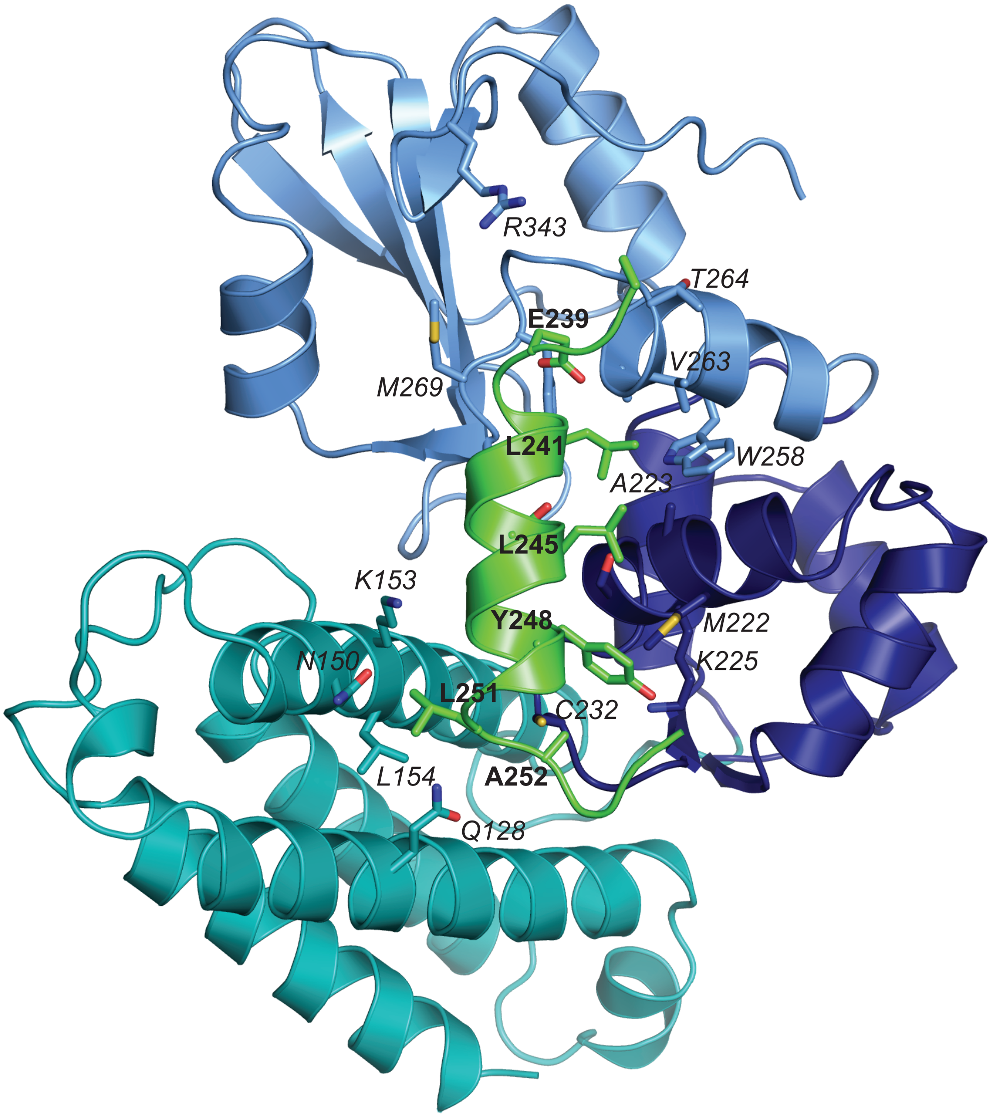

**Supplementary Figure 2**

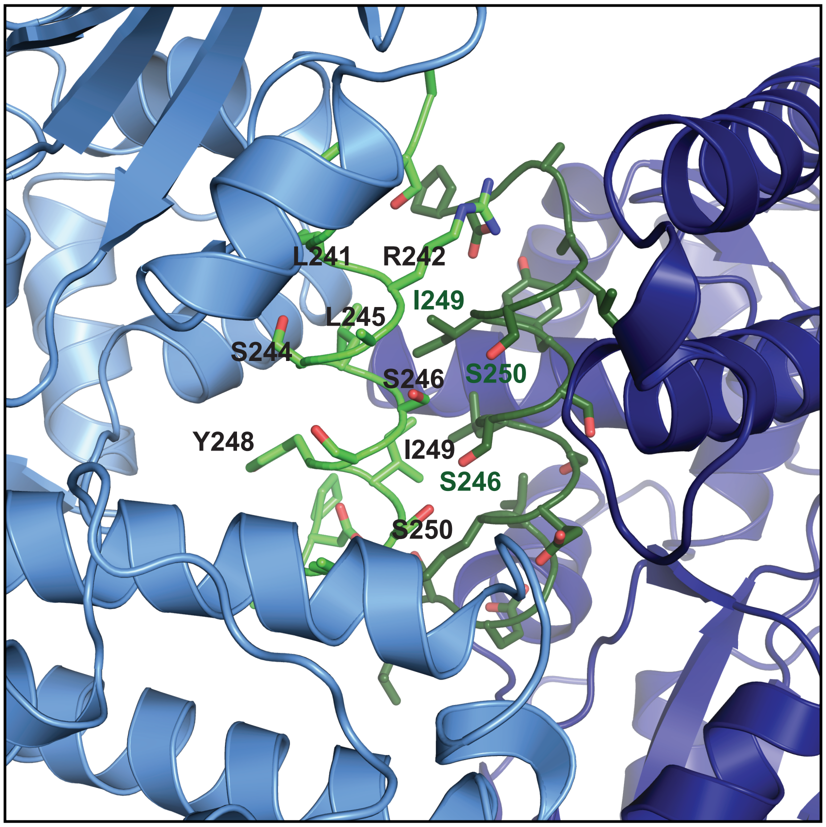

**Supplementary Figure 3**

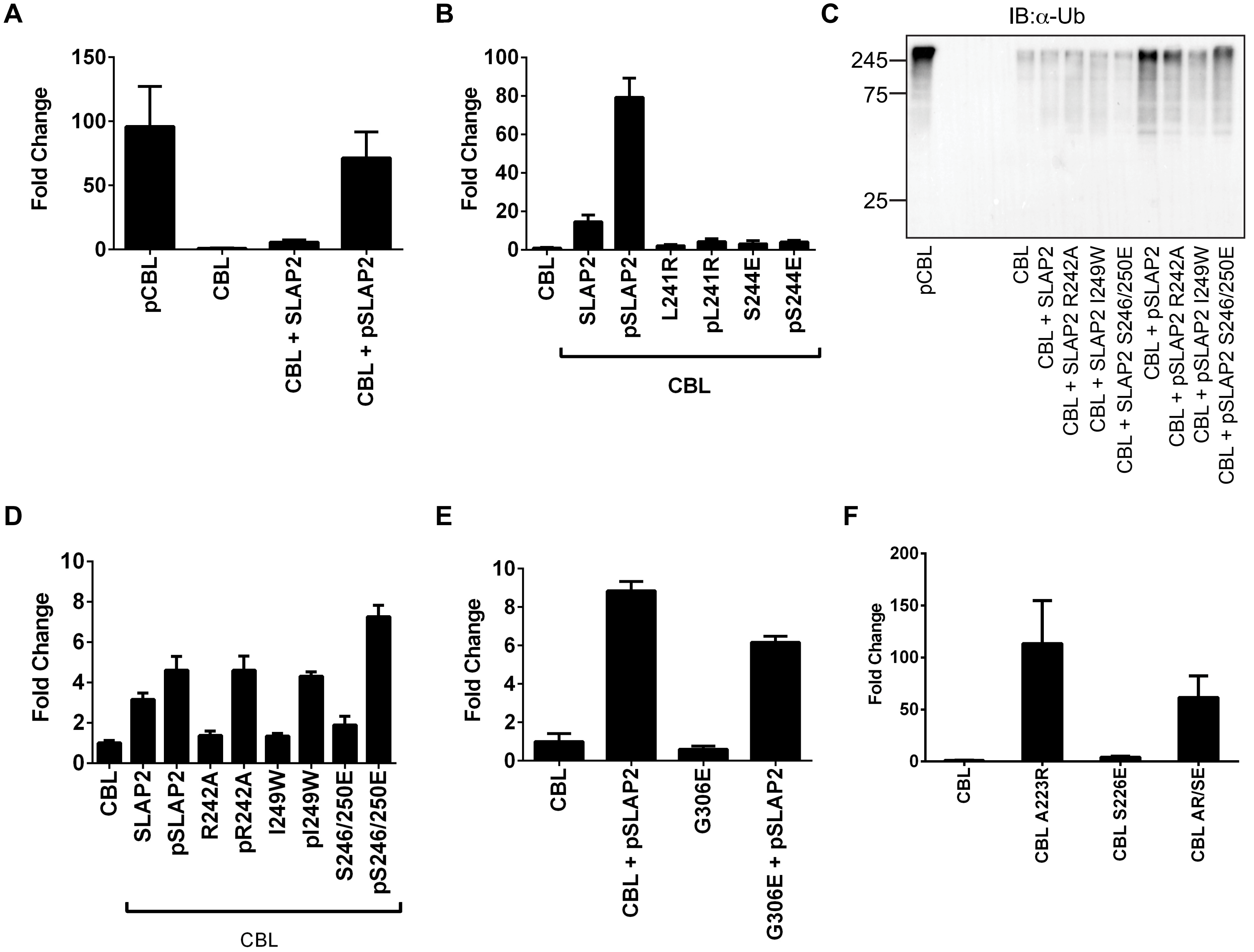

### wwPDB X-ray Structure Validation Summary Report (i)

Jun 11, 2020 – 01:40 AM EDT

| | | |
|---|---|---|
| PDB ID | : | 6XAR |
| Title | : | Structure of CBL tyrosine kinase binding domain (TKBD) with C-terminal tail of Src-like kinase protein 2 (SLAP2) |
| Deposited on | : | 2020-06-04 |
| Resolution | : | 2.50 Å(reported) |

This is a wwPDB X-ray Structure Validation Summary Report.

This report is produced by the wwPDB biocuration pipeline after annotation of the structure.

We welcome your comments at **
A user guide is available at
https://www.wwpdb.org/validation/2017/XrayValidationReportHelp
with specific help available everywhere you see the (i) symbol.

The following versions of software and data (see references (i)) were used in the production of this report:

| | | |
|---|---|---|
| MolProbity | : | 4.02b-467 |
| Xtriage (Phenix) | : | 1.13 |
| EDS | : | 2.12 |
| Percentile statistics | : | 20191225.v01 (using entries in the PDB archive December 25th 2019) |
| Refmac | : | 5.8.0158 |
| CCP4 | : | 7.0.044 (Gargrove) |
| Ideal geometry (proteins) | : | Engh & Huber (2001) |
| Ideal geometry (DNA, RNA) | : | Parkinson et al. (1996) |
| Validation Pipeline (wwPDB-VP) | : | 2.12 |

CONFIDENTIAL VALIDATION REPORT

### 1 Overall quality at a glance (i)

The following experimental techniques were used to determine the structure:
*X-RAY DIFFRACTION*

The reported resolution of this entry is 2.50 Å.

Percentile scores (ranging between 0-100) for global validation metrics of the entry are shown in the following graphic. The table shows the number of entries on which the scores are based.

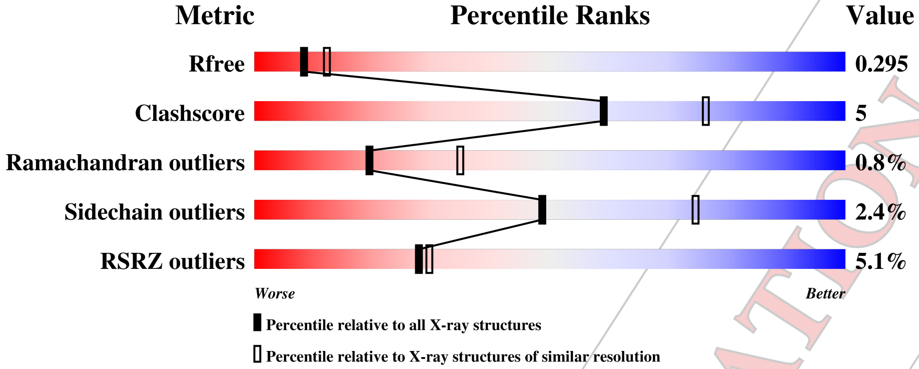


| Metric | Whole archive (#Entries) | Similar resolution (#Entries, resolution range(Å)) |
|---|---|---|
| $R_{free}$ | 130704 | 4661 (2.50-2.50) |
| Clashscore | 141614 | 5346 (2.50-2.50) |
| Ramachandran outliers | 138981 | 5231 (2.50-2.50) |
| Sidechain outliers | 138945 | 5233 (2.50-2.50) |
| RSRZ outliers | 127900 | 4559 (2.50-2.50) |

The table below summarises the geometric issues observed across the polymeric chains and their fit to the electron density. The red, orange, yellow and green segments on the lower bar indicate the fraction of residues that contain outliers for >=3, 2, 1 and 0 types of geometric quality criteria respectively. A grey segment represents the fraction of residues that are not modelled. The numeric value for each fraction is indicated below the corresponding segment, with a dot representing fractions <=5% The upper red bar (where present) indicates the fraction of residues that have poor fit to the electron density. The numeric value is given above the bar.

| Mol | Chain | Length | Quality of chain |
|---|---|---|---|
| 1 | A | 335 | 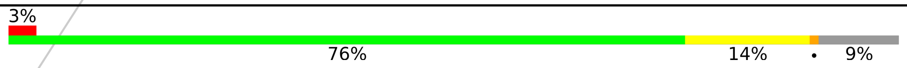 3% 76% 14% • 9% |
| 1 | B | 335 | 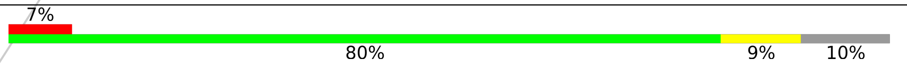 7% 80% 9% 10% |
| 2 | C | 234 | 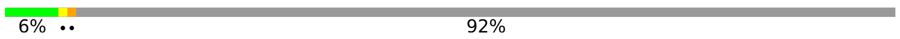 6% •• 92% |
| 2 | D | 234 | 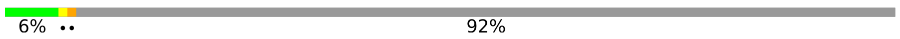 6% •• 92% |

CONFIDENTIAL VALIDATION REPORT

#### 2 Entry composition (i)

There are 4 unique types of molecules in this entry. The entry contains 5002 atoms, of which 0 are hydrogens and 0 are deuteriums.

In the tables below, the ZeroOcc column contains the number of atoms modelled with zero occupancy, the AltConf column contains the number of residues with at least one atom in alternate conformation and the Trace column contains the number of residues modelled with at most 2 atoms.

- Molecule 1 is a protein called E3 ubiquitin-protein ligase CBL.

| Mol | Chain | Residues | Atoms | ZeroOcc | AltConf | Trace |
|---|---|---|---|---|---|---|
| 1 | A | 305 | Total C N O S<br>2399 1561 399 426 13 | 0 | 0 | 0 |
| 1 | B | 300 | Total C N O S<br>2281 1487 376 408 10 | 0 | 0 | 0 |

There are 2 discrepancies between the modelled and reference sequences:

| Chain | Residue | Modelled | Actual | Comment | Reference |
|---|---|---|---|---|---|
| A | 24 | SER | GLY | conflict | UNP P22681 |
| B | 24 | SER | GLY | conflict | UNP P22681 |

- Molecule 2 is a protein called Src-like-adapter 2.

| Mol | Chain | Residues | Atoms | ZeroOcc | AltConf | Trace |
|---|---|---|---|---|---|---|
| 2 | C | 19 | Total C N O<br>140 87 22 31 | 0 | 0 | 0 |
| 2 | D | 19 | Total C N O<br>133 82 22 29 | 0 | 0 | 0 |

There are 4 discrepancies between the modelled and reference sequences:

| Chain | Residue | Modelled | Actual | Comment | Reference |
|---|---|---|---|---|---|
| C | 26 | GLY | - | expression tag | UNP Q8R4L0 |
| C | 27 | SER | - | expression tag | UNP Q8R4L0 |
| D | 26 | GLY | - | expression tag | UNP Q8R4L0 |
| D | 27 | SER | - | expression tag | UNP Q8R4L0 |

- Molecule 3 is CALCIUM ION (three-letter code: CA) (formula: Ca).

| Mol | Chain | Residues | Atoms | ZeroOcc | AltConf |
|---|---|---|---|---|---|
| 3 | B | 1 | Total Ca<br>1 1 | 0 | 0 |

*Continued on next page...*

CONFIDENTIAL VALIDATION REPORT

*Continued from previous page...*

| Mol | Chain | Residues | Atoms | ZeroOcc | AltConf |
|---|---|---|---|---|---|
| 3 | A | 1 | Total Ca<br>1 1 | 0 | 0 |

- Molecule 4 is water.

| Mol | Chain | Residues | Atoms | ZeroOcc | AltConf |
|---|---|---|---|---|---|
| 4 | A | 28 | Total O<br>28 28 | 0 | 0 |
| 4 | B | 15 | Total O<br>15 15 | 0 | 0 |
| 4 | C | 1 | Total O<br>1 1 | 0 | 0 |
| 4 | D | 3 | Total O<br>3 3 | 0 | 0 |

CONFIDENTIAL VALIDATION REPORT

### 3 Residue-property plots ⓘ

These plots are drawn for all protein, RNA and DNA chains in the entry. The first graphic for a chain summarises the proportions of the various outlier classes displayed in the second graphic. The second graphic shows the sequence view annotated by issues in geometry and electron density. Residues are color-coded according to the number of geometric quality criteria for which they contain at least one outlier: green = 0, yellow = 1, orange = 2 and red = 3 or more. A red dot above a residue indicates a poor fit to the electron density (RSRZ > 2). Stretches of 2 or more consecutive residues without any outlier are shown as a green connector. Residues present in the sample, but not in the model, are shown in grey.

• Molecule 1: E3 ubiquitin-protein ligase CBL

Chain A:

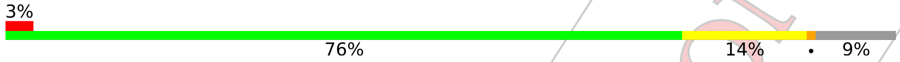


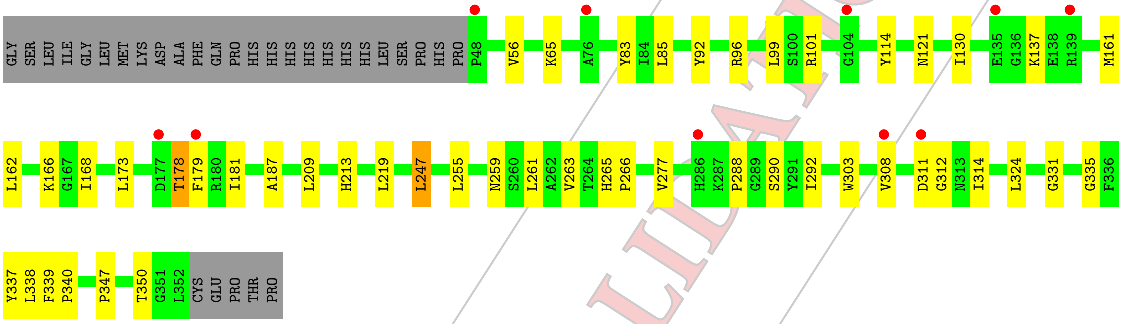


• Molecule 1: E3 ubiquitin-protein ligase CBL

Chain B:

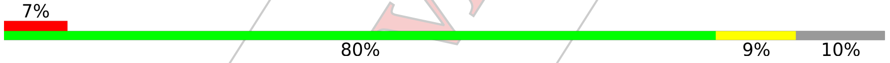


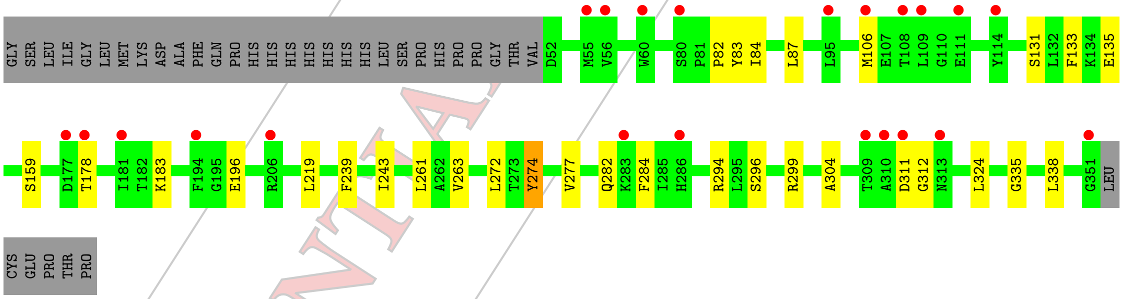


• Molecule 2: Src-like-adapter 2

Chain C:

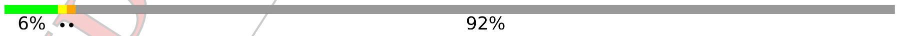


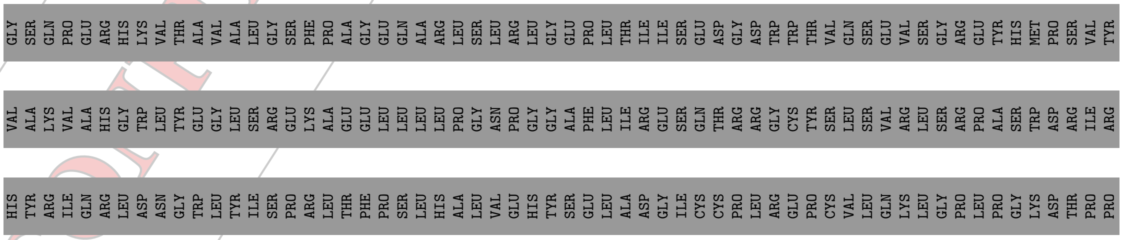


CONFIDENTIAL VALIDATION REPORT

PRO VAL THR VAL PRO THR SER SER LEU ASN TRP LYS LYS LEU ASP ARG SER LEU LEU PHE LEU GLU ALA PRO ALA SER GLY GLU ALA SER LEU L237 S238 E239 G240 L241 R242 P255 LEU ASP ASP ALA

• Molecule 2: Src-like-adapter 2

Chain D: 6% •• 92%

GLY SER GLN PRO GLU ARG HIS LYS VAL THR ALA VAL ALA LEU GLY SER PHE PRO ALA GLY GLU GLN ALA ARG LEU SER LEU ARG LEU GLY GLU PRO LEU THR ILE ILE SER GLU ASP GLY ASP TRP TRP THR VAL GLN SER GLU VAL SER GLY ARG GLU TYR HIS MET PRO SER VAL TYR

VAL ALA LYS VAL ALA HIS GLY TRP LEU TYR GLU GLY LEU SER ARG GLU LYS ALA GLU GLU LEU LEU LEU PRO GLY ASN PRO GLY GLY ALA PHE LEU ILE ARG GLU SER GLN THR ARG ARG GLY CYS TYR SER LEU SER VAL ARG LEU SER ARG PRO ALA SER TRP ASP ARG ILE ARG

HIS TYR ARG ILE GLN ARG LEU ASP ASN GLY TRP LEU TYR ILE SER PRO ARG LEU THR PHE PRO SER LEU HIS ALA LEU VAL GLU HIS TYR SER GLU LEU ALA ASP GLY ILE CYS CYS PRO LEU ARG GLU PRO CYS VAL LEU GLN LYS LEU GLY PRO LEU PRO GLY LYS ASP THR PRO PRO

PRO VAL THR VAL PRO THR SER SER LEU ASN TRP LYS LYS LEU ASP ARG SER LEU LEU PHE LEU GLU ALA PRO ALA SER GLY GLU ALA SER LEU L237 S238 E239 G240 L241 A252 P255 LEU ASP ASP ALA

CONFIDENTIAL VALIDATION REPORT

### 4 Data and refinement statistics (i)

| Property | Value | Source |
|---|---|---|
| Space group | P 1 21 1 | Depositor |
| Cell constants<br>a, b, c, $\alpha$, $\beta$, $\gamma$ | 62.73Å 86.96Å 65.26Å<br>90.00° 112.31° 90.00° | Depositor |
| Resolution (Å) | 45.32 – 2.50<br>45.32 – 2.50 | Depositor<br>EDS |
| % Data completeness<br>(in resolution range) | 83.8 (45.32-2.50)<br>83.8 (45.32-2.50) | Depositor<br>EDS |
| $R_{merge}$ | 0.08 | Depositor |
| $R_{sym}$ | (Not available) | Depositor |
| $< I/\sigma(I) >$ [1] | 2.02 (at 2.51Å) | Xtriage |
| Refinement program | PHENIX dev | Depositor |
| R, $R_{free}$ | 0.249 , 0.293<br>0.250 , 0.295 | Depositor<br>DCC |
| $R_{free}$ test set | 943 reflections (5.00%) | wwPDB-VP |
| Wilson B-factor (Å$^2$) | 40.2 | Xtriage |
| Anisotropy | 0.061 | Xtriage |
| Bulk solvent $k_{sol}$(e/Å$^3$), $B_{sol}$(Å$^2$) | 0.34 , 49.7 | EDS |
| L-test for twinning[2] | $< \|L\| >$ = 0.49, $< L^2 >$ = 0.32 | Xtriage |
| Estimated twinning fraction | 0.028 for l,-k,h | Xtriage |
| $F_o$,$F_c$ correlation | 0.90 | EDS |
| Total number of atoms | 5002 | wwPDB-VP |
| Average B, all atoms (Å$^2$) | 48.0 | wwPDB-VP |

Xtriage's analysis on translational NCS is as follows: *The largest off-origin peak in the Patterson function is 6.21% of the height of the origin peak. No significant pseudotranslation is detected.*

[1] Intensities estimated from amplitudes.

[2] Theoretical values of $< |L| >$, $< L^2 >$ for acentric reflections are 0.5, 0.333 respectively for untwinned datasets, and 0.375, 0.2 for perfectly twinned datasets.

CONFIDENTIAL VALIDATION REPORT

### 5 Model quality (i)

#### 5.1 Standard geometry (i)

Bond lengths and bond angles in the following residue types are not validated in this section: CA

The Z score for a bond length (or angle) is the number of standard deviations the observed value is removed from the expected value. A bond length (or angle) with $|Z| > 5$ is considered an outlier worth inspection. RMSZ is the root-mean-square of all Z scores of the bond lengths (or angles).

| Mol | Chain | Bond lengths | | Bond angles | |
|---|---|---|---|---|---|
| | | RMSZ | $\#\|Z\| > 5$ | RMSZ | $\#\|Z\| > 5$ |
| 1 | A | 0.21 | 0/2464 | 0.34 | 0/3345 |
| 1 | B | 0.22 | 0/2345 | 0.34 | 0/3200 |
| 2 | C | 0.25 | 0/141 | 0.46 | 0/190 |
| 2 | D | 0.20 | 0/134 | 0.36 | 0/181 |
| All | All | 0.22 | 0/5084 | 0.35 | 0/6916 |

There are no bond length outliers.

There are no bond angle outliers.

There are no chirality outliers.

There are no planarity outliers.

#### 5.2 Too-close contacts (i)

In the following table, the Non-H and H(model) columns list the number of non-hydrogen atoms and hydrogen atoms in the chain respectively. The H(added) column lists the number of hydrogen atoms added and optimized by MolProbity. The Clashes column lists the number of clashes within the asymmetric unit, whereas Symm-Clashes lists symmetry related clashes.

| Mol | Chain | Non-H | H(model) | H(added) | Clashes | Symm-Clashes |
|---|---|---|---|---|---|---|
| 1 | A | 2399 | 0 | 2326 | 28 | 1 |
| 1 | B | 2281 | 0 | 2125 | 14 | 1 |
| 2 | C | 140 | 0 | 134 | 4 | 0 |
| 2 | D | 133 | 0 | 121 | 3 | 0 |
| 3 | A | 1 | 0 | 0 | 0 | 0 |
| 3 | B | 1 | 0 | 0 | 0 | 0 |
| 4 | A | 28 | 0 | 0 | 0 | 0 |
| 4 | B | 15 | 0 | 0 | 0 | 0 |
| 4 | C | 1 | 0 | 0 | 0 | 0 |

*Continued on next page...*

CONFIDENTIAL VALIDATION REPORT

*Continued from previous page...*

| Mol | Chain | Non-H | H(model) | H(added) | Clashes | Symm-Clashes |
|---|---|---|---|---|---|---|
| 4 | D | 3 | 0 | 0 | 0 | 0 |
| All | All | 5002 | 0 | 4706 | 45 | 1 |

The all-atom clashscore is defined as the number of clashes found per 1000 atoms (including hydrogen atoms). The all-atom clashscore for this structure is 5.

The worst 5 of 45 close contacts within the same asymmetric unit are listed below, sorted by their clash magnitude.

| Atom-1 | Atom-2 | Interatomic distance (Å) | Clash overlap (Å) |
|---|---|---|---|
| 1:A:308:VAL:HG12 | 1:A:314:ILE:HG12 | 1.76 | 0.68 |
| 1:A:335:GLY:HA2 | 1:A:338:LEU:HD21 | 1.82 | 0.62 |
| 1:A:219:LEU:HD23 | 1:B:219:LEU:HD23 | 1.81 | 0.62 |
| 1:A:178:THR:OG1 | 1:A:178:THR:O | 2.16 | 0.61 |
| 1:B:311:ASP:N | 1:B:312:GLY:HA2 | 2.16 | 0.59 |

All (1) symmetry-related close contacts are listed below. The label for Atom-2 includes the symmetry operator and encoded unit-cell translations to be applied.

| Atom-1 | Atom-2 | Interatomic distance (Å) | Clash overlap (Å) |
|---|---|---|---|
| 1:A:178:THR:OG1 | 1:B:274:TYR:OH[1_556] | 2.18 | 0.02 |

#### 5.3 Torsion angles (i)

##### 5.3.1 Protein backbone (i)

In the following table, the Percentiles column shows the percent Ramachandran outliers of the chain as a percentile score with respect to all X-ray entries followed by that with respect to entries of similar resolution.

The Analysed column shows the number of residues for which the backbone conformation was analysed, and the total number of residues.

| Mol | Chain | Analysed | Favoured | Allowed | Outliers | Percentiles | |
|---|---|---|---|---|---|---|---|
| 1 | A | 303/335 (90%) | 286 (94%) | 15 (5%) | 2 (1%) | 22 | 39 |
| 1 | B | 298/335 (89%) | 278 (93%) | 18 (6%) | 2 (1%) | 22 | 39 |
| 2 | C | 17/234 (7%) | 15 (88%) | 2 (12%) | 0 | 100 | 100 |
| 2 | D | 17/234 (7%) | 15 (88%) | 1 (6%) | 1 (6%) | 1 | 1 |
| All | All | 635/1138 (56%) | 594 (94%) | 36 (6%) | 5 (1%) | 19 | 35 |

CONFIDENTIAL VALIDATION REPORT

All (5) Ramachandran outliers are listed below:

| Mol | Chain | Res | Type |
|---|---|---|---|
| 1 | A | 137 | LYS |
| 2 | D | 239 | GLU |
| 1 | A | 350 | THR |
| 1 | B | 263 | VAL |
| 1 | B | 196 | GLU |

##### 5.3.2 Protein sidechains (i)

In the following table, the Percentiles column shows the percent sidechain outliers of the chain as a percentile score with respect to all X-ray entries followed by that with respect to entries of similar resolution.

The Analysed column shows the number of residues for which the sidechain conformation was analysed, and the total number of residues.

| Mol | Chain | Analysed | Rotameric | Outliers | Percentiles | |
|---|---|---|---|---|---|---|
| 1 | A | 249/299 (83%) | 247 (99%) | 2 (1%) | 81 | 93 |
| 1 | B | 225/299 (75%) | 219 (97%) | 6 (3%) | 44 | 71 |
| 2 | C | 16/200 (8%) | 14 (88%) | 2 (12%) | 4 | 8 |
| 2 | D | 14/200 (7%) | 12 (86%) | 2 (14%) | 3 | 6 |
| All | All | 504/998 (50%) | 492 (98%) | 12 (2%) | 49 | 74 |

5 of 12 residues with a non-rotameric sidechain are listed below:

| Mol | Chain | Res | Type |
|---|---|---|---|
| 1 | B | 178 | THR |
| 1 | B | 183 | LYS |
| 2 | C | 241 | LEU |
| 1 | B | 133 | PHE |
| 2 | C | 239 | GLU |

Some sidechains can be flipped to improve hydrogen bonding and reduce clashes. There are no such sidechains identified.

##### 5.3.3 RNA (i)

There are no RNA molecules in this entry.

CONFIDENTIAL VALIDATION REPORT

#### 5.4 Non-standard residues in protein, DNA, RNA chains (i)

There are no non-standard protein/DNA/RNA residues in this entry.

#### 5.5 Carbohydrates (i)

There are no carbohydrates in this entry.

#### 5.6 Ligand geometry (i)

Of 2 ligands modelled in this entry, 2 are monoatomic - leaving 0 for Mogul analysis.

There are no bond length outliers.

There are no bond angle outliers.

There are no chirality outliers.

There are no torsion outliers.

There are no ring outliers.

No monomer is involved in short contacts.

#### 5.7 Other polymers (i)

There are no such residues in this entry.

#### 5.8 Polymer linkage issues (i)

There are no chain breaks in this entry.

CONFIDENTIAL VALIDATION REPORT

### 6 Fit of model and data (i)

#### 6.1 Protein, DNA and RNA chains (i)

In the following table, the column labelled '#RSRZ> 2' contains the number (and percentage) of RSRZ outliers, followed by percent RSRZ outliers for the chain as percentile scores relative to all X-ray entries and entries of similar resolution. The OWAB column contains the minimum, median, $95^{th}$ percentile and maximum values of the occupancy-weighted average B-factor per residue. The column labelled 'Q< 0.9' lists the number of (and percentage) of residues with an average occupancy less than 0.9.

| Mol | Chain | Analysed | <RSRZ> | #RSRZ>2 | | | OWAB(Å$^2$) | Q<0.9 |
|---|---|---|---|---|---|---|---|---|
| 1 | A | 305/335 (91%) | 0.23 | 10 (3%) | 46 | 50 | 24, 42, 71, 81 | 0 |
| 1 | B | 300/335 (89%) | 0.59 | 22 (7%) | 15 | 15 | 26, 57, 80, 92 | 0 |
| 2 | C | 19/234 (8%) | 0.36 | 0 | 100 | 100 | 30, 36, 51, 57 | 0 |
| 2 | D | 19/234 (8%) | 0.64 | 1 (5%) | 26 | 28 | 26, 34, 56, 63 | 0 |
| All | All | 643/1138 (56%) | 0.42 | 33 (5%) | 28 | 29 | 24, 47, 76, 92 | 0 |

The worst 5 of 33 RSRZ outliers are listed below:

| Mol | Chain | Res | Type | RSRZ |
|---|---|---|---|---|
| 1 | A | 177 | ASP | 6.0 |
| 1 | B | 108 | THR | 5.7 |
| 1 | B | 309 | THR | 4.9 |
| 1 | B | 106 | MET | 4.1 |
| 1 | B | 56 | VAL | 3.5 |

#### 6.2 Non-standard residues in protein, DNA, RNA chains (i)

There are no non-standard protein/DNA/RNA residues in this entry.

#### 6.3 Carbohydrates (i)

There are no carbohydrates in this entry.

#### 6.4 Ligands (i)

In the following table, the Atoms column lists the number of modelled atoms in the group and the number defined in the chemical component dictionary. The B-factors column lists the minimum, median, $95^{th}$ percentile and maximum values of B factors of atoms in the group. The column labelled 'Q< 0.9' lists the number of atoms with occupancy less than 0.9.

CONFIDENTIAL VALIDATION REPORT

| Mol | Type | Chain | Res | Atoms | RSCC | RSR | B-factors(Å$^2$) | Q<0.9 |
|---|---|---|---|---|---|---|---|---|
| 3 | CA | B | 401 | 1/1 | 0.90 | 0.07 | 50,50,50,50 | 0 |
| 3 | CA | A | 401 | 1/1 | 0.94 | 0.09 | 48,48,48,48 | 0 |

#### 6.5 Other polymers ⓘ

There are no such residues in this entry.

CONFIDENTIAL VALIDATION REPORT
